# Spatial metabolomics reveal divergent cardenolide processing in the monarch butterfly (*Danaus plexippus*) and the common crow (*Euploea core*)

**DOI:** 10.1101/2022.06.02.494161

**Authors:** Domenic Dreisbach, Dhaka R. Bhandari, Anja Betz, Linda Tenbusch, Andreas Vilcinskas, Bernhard Spengler, Georg Petschenka

## Abstract

Although being famous for sequestering milkweed cardenolides, the mechanism of sequestration and where cardenolides are localized in caterpillars of the monarch butterfly (*Danaus plexippus*) is still unknown. While monarchs tolerate cardenolides by a resistant Na^+^/K^+^-ATPase, it is unclear how closely related species such as the non-sequestering common crow (*Euploea core*) cope with these toxins. Using novel atmospheric-pressure scanning microprobe matrix-assisted laser/desorption ionization mass spectrometry imaging, we compared the distribution of cardenolides in caterpillars of *D. plexippus* and *E. core*. Specifically, we tested at which physiological scale quantitative differences between both species are mediated and how cardenolides distribute across body tissues. Whereas *D. plexippus* sequestered most cardenolides from milkweed (*Asclepias curassavica*), no cardenolides were found in the tissues of *E. core*. Remarkably, quantitative differences already manifest in the gut lumen: while monarchs retain and accumulate cardenolides above plant concentrations, the toxins are degraded in the gut lumen of crows. We visualized cardenolide transport over the monarch midgut epithelium and identified integument cells as the final site of storage where defenses might be perceived by predators. Our study provides molecular insight into cardenolide sequestration and highlights the great potential of mass spectrometry imaging for understanding the kinetics of multiple compounds including endogenous metabolites, plant toxins, or insecticides in insects.

## Introduction

As a product of reciprocal coevolution, plants possess a plethora of defenses against herbivorous insects and other antagonists. These traits include the production of toxic secondary plant metabolites, which protect plants by affecting herbivores directly (Dussourd and Hoyle, 2000; Narberhaus et al., 2005), impair their growth and development (Ayres et al., 1997; Cresswell et al., 1992), or lower the digestibility of the plant diet (Fraenkel, 1959; Koiwa et al., 1997). Remarkably, many insects are not only able to cope with a toxic diet, but also sequester (i.e. accumulate and store) plant toxins in their body tissues to defend themselves against predators and parasitoids (Beran and Petschenka, 2022; Duffey, 1980; Opitz and Müller, 2009). Although sequestration is a widespread phenomenon among herbivorous insects including several important pests (Beran et al., 2014; Kazana et al., 2007; Robert et al., 2017; Yang et al., 2021), the underlying physiological mechanisms and especially the transport of toxins across the gut epithelium or the spatial distribution of plant toxins across insect body tissues are largely unknown.

The monarch butterfly (*Danaus plexippus*) is an important model for insect-plant coevolution (Agrawal et al., 2021; Karageorgi et al., 2019; Petschenka and Agrawal, 2015) and is famous for the sequestration of toxic cardenolides from its host plant milkweed (*Asclepias* spp., Apocynaceae) (Brower et al., 1968; Frick and Wink, 1995; Parsons, 1965; Reichstein et al., 1968). Cardenolides are potent inhibitors of Na^+^/K^+^-ATPase (Agrawal et al., 2012; Schatzmann, 1953), an essential cation carrier ubiquitously expressed in animal cells. Remarkably, monarch larvae sequester these toxins in high amounts and can tolerate them by means of a modified Na^+^/K^+^-ATPase, carrying a few amino acid substitutions in the cardenolide binding site of the enzyme (Dobler et al., 2012; Holzinger and Wink, 1996; Vaughan and Jungreis, 1977).

Despite feeding on cardenolide producing plants as well, the closely-related milkweed butterfly *Euploea core* possesses a non-resistant Na^+^/K^+^-ATPase and does not sequester cardenolides (Malcom and Rothschild, 1983; Petschenka and Agrawal, 2015). Although it is not yet understood how caterpillars of *E. core* cope with dietary cardenolides, tolerance is potentially mediated on the level of the caterpillars’ gut and may involve degradation of the compounds (Petschenka and Agrawal, 2015) or epithelial barriers (Dobler et al., 2015). Using a comparative approach involving caterpillars of both species raised on the same species of milkweed, we set out to test on which physiological scale the differences in toxin distribution are mediated.

Cardenolides consist of a 23 C four-ring steroid skeleton bearing a five-membered lactone ring bound to C17 and are frequently attached to a sugar moiety. Approximately 200 cardenolides are known from various milkweed species (Züst et al., 2019), and up to 21 different cardenolides were identified in *Asclepias curassavica* (Zhang et al., 2014), one of the two most important host plants of the monarch butterfly (Agrawal et al., 2021). Although sequestration in monarch butterflies was already described in the 1960s (Brower et al., 1968; Reichstein et al., 1968), it is still largely unknown how sequestration is mediated mechanistically and where cardenolides are localized in the caterpillar body. However, a comprehensive view on the spatial distribution and abundance of cardenolides across caterpillar body tissues is a prerequisite to understand the molecular mechanisms of sequestration and the function of stored defenses in ecological interactions with predators and parasitoids. Along these lines, comparisons to closely related but non-sequestering species such as *E. core*, are mandatory to reveal specific adaptations related to cardenolide sequestration in the monarch butterfly.

Among MSI methods, atmospheric-pressure scanning-microprobe matrix-assisted laser desorption/ionization (AP-SMALDI) MSI is one of the most advanced techniques regarding spatial resolution and sensitivity and therefore is of special interest for the spatial metabolomic characterization of complex molecular systems in the fields of chemistry, biology and medicine (Koestler et al., 2008; Kompauer et al., 2017a, 2017b; Spengler et al., 1994; Spengler and Hubert, 2002). For example, metabolite, lipid and peptide distributions were successfully visualized with spatial resolutions of up to 5 μm on various biological surfaces and within native organisms, thereby linking molecular structures to biological functions and origins (Geier et al., 2021, 2020; Iwama et al., 2021; Kadesch et al., 2020b, 2020a, 2019; Kompauer et al., 2017b).

In contrast to vacuum-ionization MSI methods such as conventional MALDI and secondary ion mass spectrometry (SIMS), AP-SMALDI MSI allows experiments to be performed under near-physiological conditions (i.e. physiological temperature and pressure). Therefore, AP-SMALDI MSI overcomes vacuum-induced sample damage, loss of volatile compounds and does not require sample fixation and/or sample drying. The most recent instrumental approach in AP-SMALDI MSI eliminates height-related measurement artifacts by utilizing laser triangulation for pixel-wise autofocusing, thereby enabling the spatially-resolved molecular analysis of delicate and non-planar sample surfaces, such as the fragile epithelium of the caterpillar integument and heterogeneous tissue types of insects. To this date, the experimental bottlenecks described above had limited the application of high-resolution MSI in the field of insect research (Yang et al., 2020).

Taking advantage of recent AP-SMALDI MSI developments, we here set out to study the distribution of ingested and sequestered cardenolides within their native, spatial context in caterpillars of *D. plexippus* and *E. core*. We followed an MSI-based spatial metabolomics approach to study the fate of 10 cardenolides in tissue sections of both species at high lateral resolution, thus providing novel molecular insight into the process of cardenolide sequestration in *D. plexippus* and cardenolide tolerance in *E. core*. In contrast to previous work on cardenolide sequestration in monarch caterpillars, our study provides detailed information on cardenolide structures, selectivity of absorption, and most importantly, the distribution of individual cardenolides across different tissues involved in the process of sequestration such as the midgut epithelium and the caterpillar integument. While cardenolides are degraded in the gut lumen of *E. core* caterpillars, they accumulate in the gut lumen of monarch caterpillars representing a novel mechanism of processing plant toxins in herbivorous insects.

Besides providing insight into the process of sequestration in the monarch butterfly and revealing mechanistic differences between two closely related caterpillars coping with the same class of plant toxins but having divergent ecological strategies, our study highlights the potential of high-resolution AP-SMALDI MSI emerging as a powerful platform for researchers to in-situ visualize the distribution and kinetics of metabolites in the spatial context of insect tissues and cells, with applications ranging from chemically-mediated plant-insect interactions to insecticide research.

## Results

### Spatial distribution of cardenolides in *D. plexippus* and *E. core*

AP-SMALDI MSI experiments (20 μm and 35 μm step size) conducted on transversal caterpillar sections (Figs. 1 and S1) revealed the presence of cardenolides throughout the body tissues of *D. plexippus* caterpillars (Fig. 1b) including the midgut epithelium, the body cavity (fat body and hemolymph), and especially the integument from where we obtained the strongest cardenolide signals. In contrast, cardenolides were entirely restricted to the gut lumen in caterpillars of *E. core* and were absent from its body tissues. Remarkably, cardenolides appeared at much higher intensities in guts of *D. plexippus* compared to guts of *E. core* when images were adjusted to the same scale. This pattern was not restricted to the isomers calotropin/calactin but also apparent for uscharidin and voruscharin (Figs. S1 and S2, not all cardenolides were compared). In total, we found 10 different cardenolides in both caterpillar species and leaves of *A. curassavica* (Tab. 1 and Fig. S3-7).

**Figure 1.**
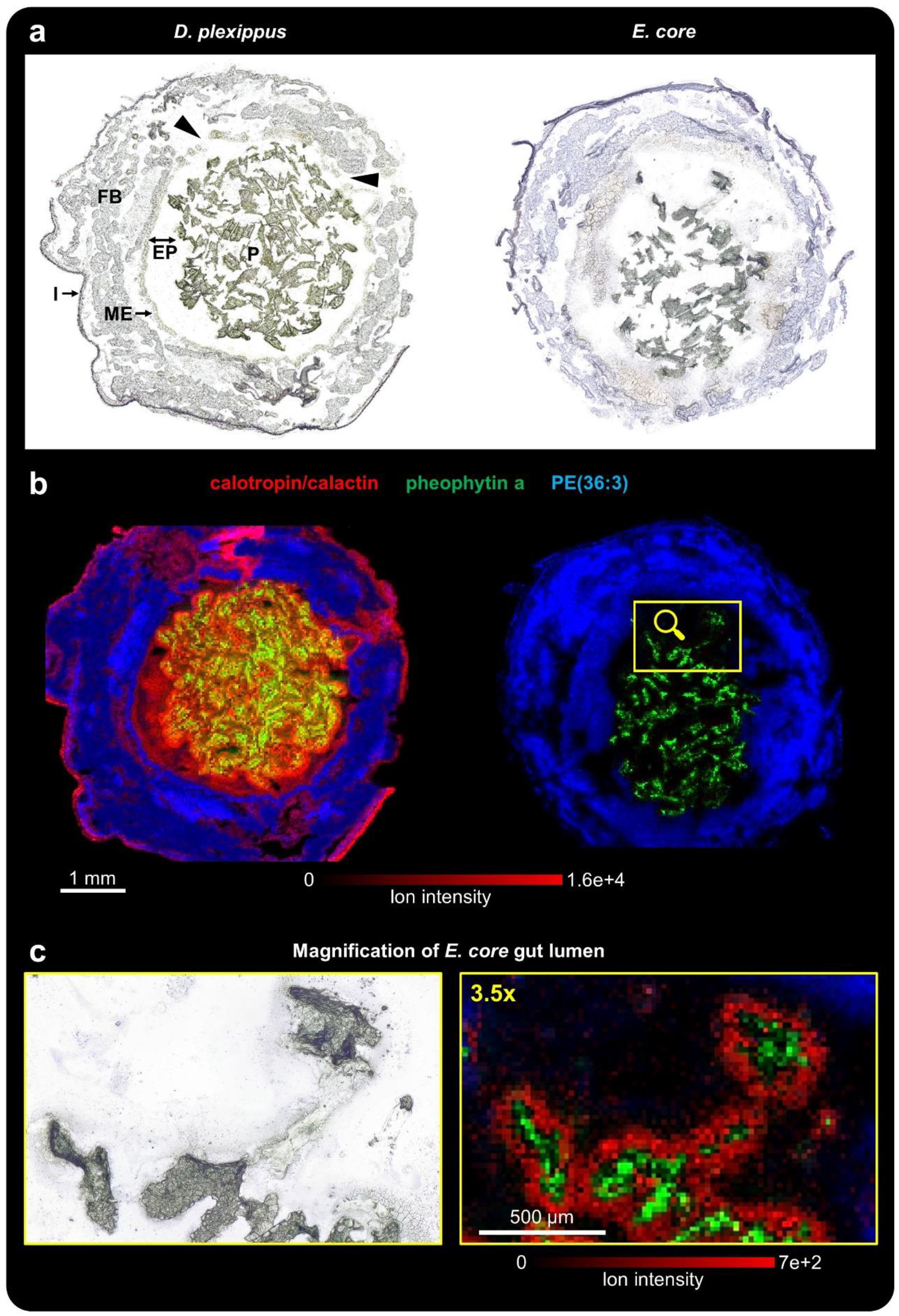
AP-SMALDI MSI of transversal sections of caterpillars of *D. plexippus* (biological replicate 1) and *E. core* (biological replicate 1). (**a**) Optical image of the transversal section of a fifth instar caterpillar of *D. plexippus* (left) and *E. core* (right) before matrix application. P: plant material, EP: ectoperitrophic space, ME: midgut epithelium, FB: fat body, I: integument. (**b**) Corresponding RGB overlay images obtained with 35 μm (*D. plexippus*) and 20 μm (*E. core*) step size, showing the spatial distribution of the cardenolide calotropin and/or its isomer calactin ([M+K]^+^, red) at *m/z* 571.2304, the chlorophyll derivative pheophytin a at *m/z* 909.5288 ([M+K]^+^, green) as a chemical marker for plant tissue, and the animal lipid PE(36:3) ([M+K]^+^, blue) as a chemical marker for animal tissues. Both RGB overlay images are normalized to the same intensity scale. The *D. plexippus* gut epithelium was damaged at two areas (highlighted in the optical image), causing potential analyte delocalization in the corresponding hemolymph area. (**c**) Magnification of the optical image and RGB overlay image for the highlighted area of the *E. core* gut lumen. For this ion image, the intensity scale of calotropin/calactin was adjusted to highlight the cardenolide distribution at the fringes of leaf pieces in the gut lumen of *E. core*.

Notably, cardenolides (Fig. 1b at the example of [calotropin/calactin+K]^+^) were visible across the entire gut lumen of *D. plexippus* including the ectoperitrophic space (i.e. the region between the peritrophic envelope surrounding the food bolus and the midgut epithelium). In midguts of *E. core*, in contrast, they were exclusively detected at the fringes of *A. curassavica* leaf pieces (Figs. 1c and S1b) and were absent in the liquid phase of the gut lumen. In addition, to transversal sections, we carried out whole-body MSI (45 μm step size) on longitudinal sections of final instar caterpillars and found according patterns of cardenolide distributions along the entire gut passage (Figs. 2 and S8).

**Figure 2.**
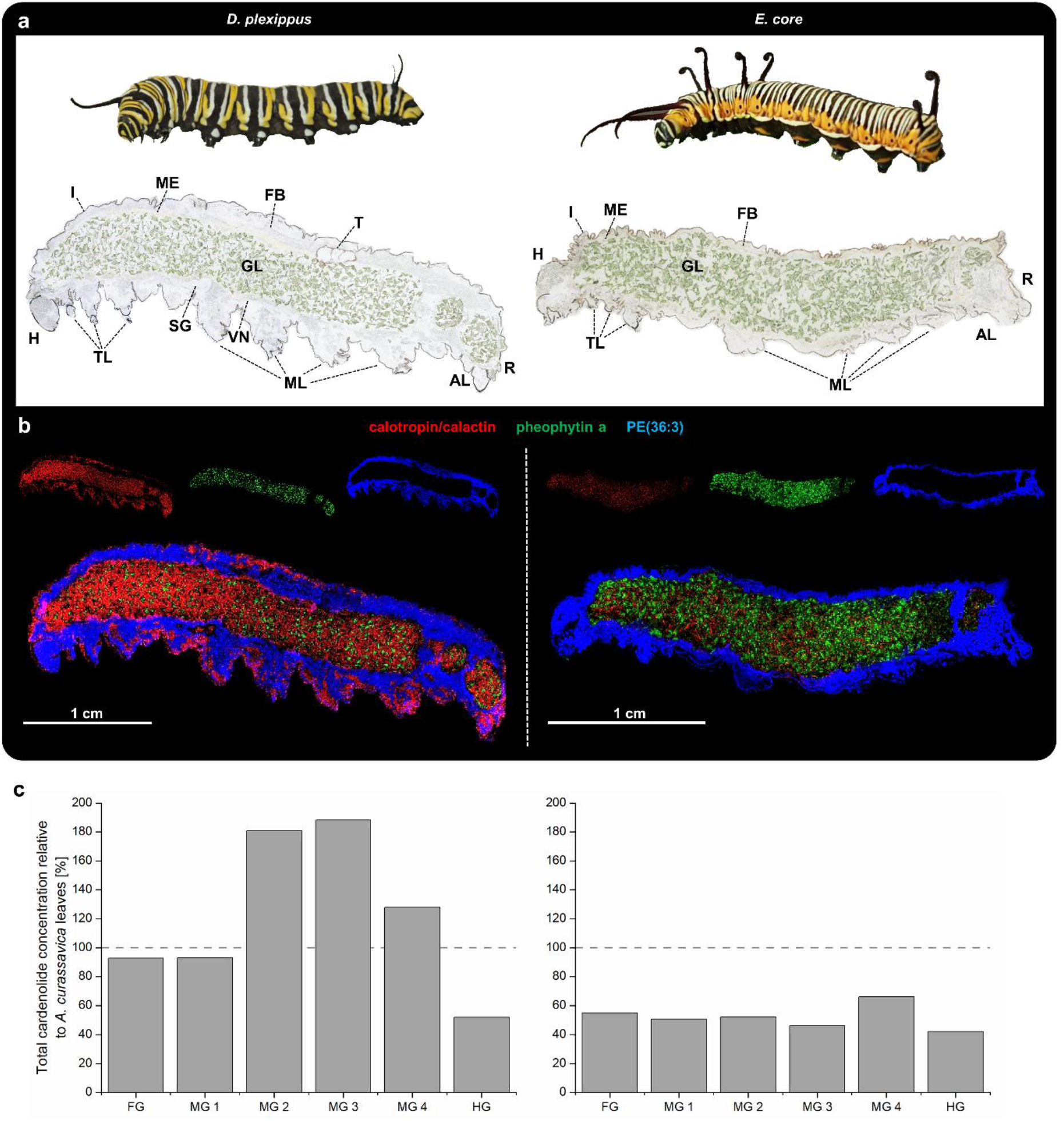
Whole-body AP-SMALDI MSI of fifth instar *D. plexippus* (biological replicate 3) and *E. core* caterpillars (biological replicate 3). (**a**) Optical image showing live caterpillars and longitudinal *D. plexippus* (left) and *E. core* (right) sections before matrix application. H: head, I: integument, TL: thoracic legs, SG: probably salivary gland, VN: probably ventral nerve code, ML: abdominal prolegs, AL: anal prolegs, R: rectum, T: testis, GL: gut lumen, FB: fat body, ME: midgut epithelium. (**b**) Corresponding RGB overlay images obtained with 45 μm step size, showing the spatial distribution of calactin/calotropin ([M+K]^+^, at *m/z* 571.2304 (red), pheophytin a ([M+K]^+^, green) at *m/z* 909.5292 to highlight ingested *A. curassavica* plant material, and PE(36:3) ([M+K]^+^, blue) at *m/z* 780.4941 serving as a marker for insect tissue, such as gut epithelium and fat body. Both RGB overlay images are normalized to the same intensity scale. (**c**) Schematic representation of total cardenolide concentrations across the gut passage in caterpillars of *D. plexippus* and *E. core* relative to the total cardenolide concentration in *A. curassavica* leaves based on means of six (*E. core*) and nine (*D. plexippus*) dissected caterpillars and milkweed leaves. Horizontal lines indicate the cardenolide concentration in plant material (i.e. 100%). Please see Fig. S12 for a plot based on absolute quantification data.

We compared relative quantities of the five most abundant cardenolides (the indistinguishable isomers calotropin/calactin counted as one) based on MSI data *in silico*. For this purpose, we selected seven comparable regions of interest (ROI) in transversal and longitudinal sections of *D. plexippus* and *E. core* caterpillars and analyzed cardenolide intensities at the fringes of leaf pieces in the gut. According to the visual differences apparent in Fig.1 and 2, the *in silico* analysis revealed that calotropin/calactin was 6.8 x, calotoxin 2.1 x, frugoside 2.6 x, and uscharidin 1.8 x more abundant in *D. plexippus* compared to *E. core* (p < 0.001, p = 0.031, p = 0.005, p = 0.02, no difference for asclepin: 1.15 x, p = 0.2; n = 4, i.e. two transversal and two longitudinal sections from individual caterpillars per species; all t-tests assuming unequal variances, Figure S9). Notably, the analyzed regions of interest (see methods and Fig. S9) included leaf particles and their fringes where cardenolides were detectable. More distantly from the leaf fragments, the difference between the liquid phase of *D. plexippus* and *E. core* would certainly have been even more pronounced.

Besides local quantitative differences between the two caterpillar species, our total cardenolide estimate based on HPLC-DAD analyses (i.e. all cardenolide peaks integrated) of dissected freeze-dried gut regions (foregut, four segments of the midgut and hindgut) revealed a heterogeneous quantitative distribution of cardenolides along the gut passage in *D. plexippus* but not in *E. core*. Total cardenolide concentrations were constant across all gut regions of *E. core* caterpillars (foregut: FG, four sequential midgut portions: MG1-4, and hindgut: HG; F_5,25_ = 1.044, p = 0.414, n = 6; Figs. 12 for HPLC-DAD and S13 for HPLC-MS). Contrastingly, in caterpillars of *D. plexippus*, cardenolide concentrations across all gut regions differed substantially (F_5,38_ = 8.312, p < 0.001; n = 7-9, see legend of Fig. S12) and the cardenolide concentration in the hindgut was lower compared to all other gut regions except of foregut and midgut region 1 (FG vs. HG: p = 0.577; MG1 vs. HG: p = 0.085; MG2-4 vs. HG: p < 0.001; Tukey HSD; Fig. S20). Concentrations of gut cardenolides differed within caterpillar individuals in both species (*E. core:* F_5,25_ = 22.015, p < 0.001; *D. plexippus:* F_8,38_ = 2.352, p < 0.037).

When comparing total cardenolide concentrations across midgut portions (i.e. foregut and hindgut excluded) of monarch caterpillars and leaves of *A. curassavica*, we found higher cardenolide concentrations compared to milkweed leaves in midgut portions 2 and 3 (F_4,32_ = 5.622, p = 0.002) and a trend of cardenolide accumulation in the central region of the midgut with concentrations of midgut portion 2 and 3 being twice as high compared to midgut portion 1 (F_3,24_ = 5.121, p = 0.007; MG1 vs. MG2: p = 0.018; midgut 1 vs. midgut 3: p = 0.01; Tukey HSD; Fig. S12). In contrast, cardenolide concentrations in gut portions of *E. core* were constantly lower than in plant material except for midgut portion 4 (plant vs. FG, MG1, MG2, MG3, MG4, HG: p = 0.046; 0.023; 0.029; 0.01; 0.228; 0.005; Tukey HSD). Relatively constant toxin concentrations across guts of *E. core* and a heterogeneous distribution in *D. plexippus* were also observed based on absolute quantification of individual cardenolides via HPLC-MS (Fig. S13).

Notably, the ratio of the stereoisomers calotropin and calactin in the gut passage differed substantially between both caterpillar species. Across the whole gut passage of *D. plexippus*, ratios of calotropin and calactin concentrations were rather constant (calotropin: calactin = 1.35; t-test, p = 0.15; Figure S14). Contrastingly, in the gut material of *E. core*, the concentration of calotropin had an 8.15 x higher concentration relative to calactin (t-test, p < 0.001) (Figure S14). Regionally, the largest difference in the calotropin/calactin ratio between both species was determined in the first portion of the midgut (Figure S15). For comparison, we also analyzed the content of both stereoisomers in leaf tissue of the host plant *A. curassavica* (1.61-fold for calotropin relative to calactin, t-test, p = 0.05) and for integument tissue of *D. plexippus* (4.50-fold of calotropin relative to calactin, t-test, p < 0.001; n = 3 for all comparisons) (Figure S15).

### Retention of cardenolides in the gut lumen of *D. plexippus*

Based on our observation on cardenolide accumulation in guts of *D. plexippus*, we tested the hypothesis that cardenolides are retained in the midgut lumen of *D. plexippus* caterpillars. For this purpose, we analyzed last instar caterpillars of *D. plexippus* which were raised on *A. curassavica* but were fed with the cardenolide-free plant *Oxypetalum coeruleum* (Warashina and Shirota, 2021) for three hours before sampling using AP-SMALDI MSI. Remarkably, even after several rounds of purging with cardenolide-free *O. coeruleum*, we still detected cardenolides in the gut lumen. The preferentially-sequestered cardenolides calotropin/calactin (Fig. 3b), calotoxin (Fig. 3c), calotropagenin (Fig. S16a) and frugoside (Fig. S16b), as well as uscharidin (Fig. S16c), which is the dominant cardenolide in *A. curassavica* leaves, were still present in the interstices between the *O. coeruleum* leaf particles in the gut although at lower abundance compared to a caterpillar freshly harvested from *A. curassavica* (see Fig. 1, Fig. S17; representative MSI-based relative quantification between biological replicate 1 and biological replicate 5 showed 2.7 x, 1.1 x, 2.1 x reduction for calotropin/calactin, calotoxin, and frugoside, respectively; t-test, p < 0.001 for all comparisons; see Fig. S17 legend for details regarding *in silico* quantification). Notably, the dominant *A. curassavica* leaf cardenolides, voruscharin, uscharin and asclepin were not detected anymore. Thus, our data suggest that sequestered cardenolides are retained in the gut lumen and are not moving linearly along with the gut contents.

**Figure 3.**
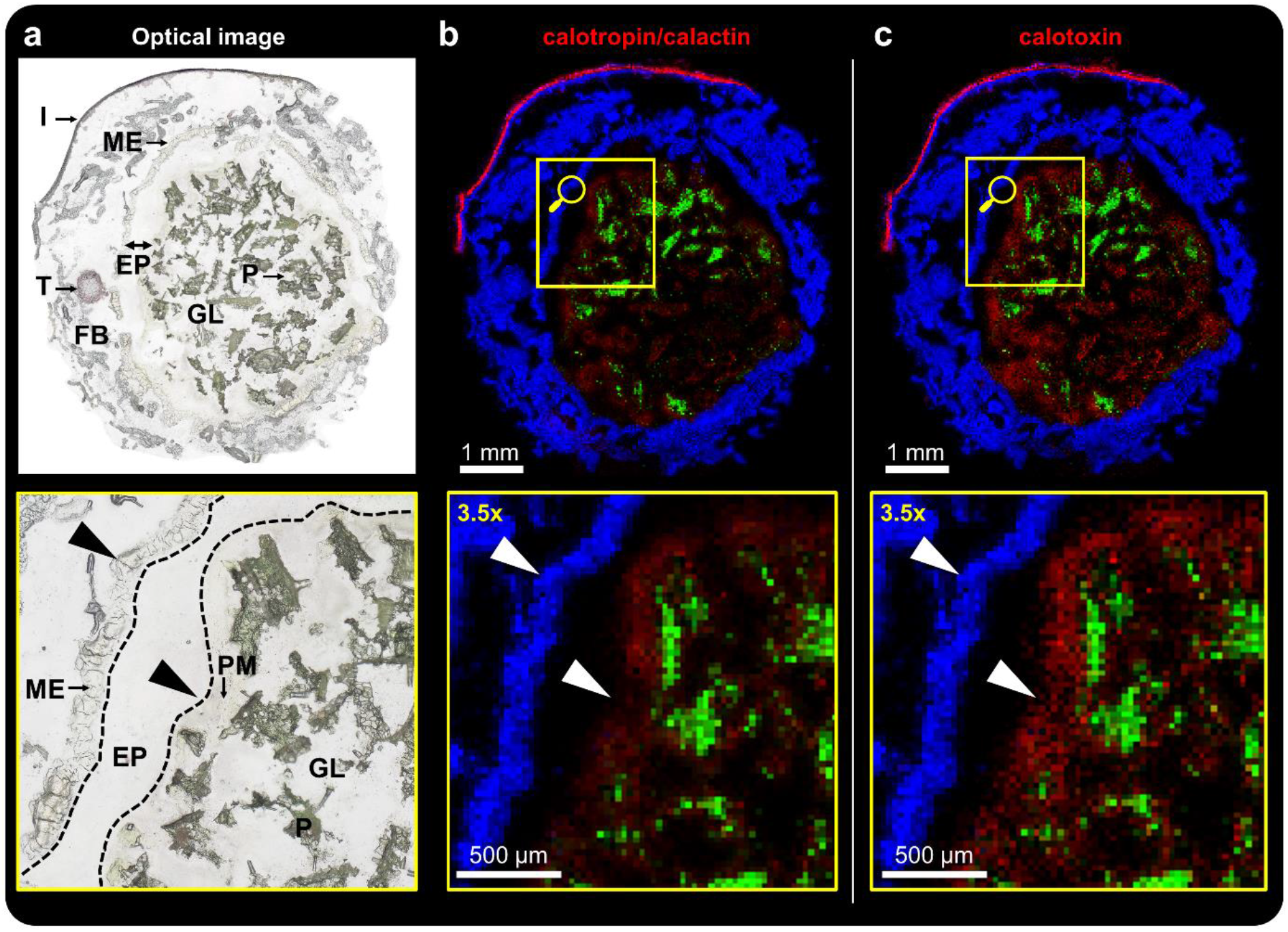
AP-SMALDI MSI (25 μm step size) of *D. plexippus* (biological replicate 5), fed with the non-toxic plant *Oxypetalum coeruleum* for 3 hours before sampling. (**a**) Optical image of transversal *D. plexippus* section and magnified view of the outlined region. P: *O. coeruleum* plant material, EP: ectoperitrophic space, PM: peritrophic matrix, ME: midgut epithelium, T: testis, FB: fat body, I: integument. (**b,c**) Corresponding RGB overlay images and magnified views of the outlined region in **a**, showing (**b**) calactin/calotropin ([M+K]^+^, red) at *m/z* 571.2304 and (**c**) calotoxin ([M+K]^+^, red) at *m/z* 587.2251 and (**b,c**) pheophytin a ([M+K]^+^) at *m/z* 909.5292 as a chemical marker for plant tissue in green and PE(36:3) ([M+K]^+^) at *m/z* 780.4942 as a chemical marker for insect tissue is shown in blue.

In contrast to monarch caterpillars that were analyzed directly after feeding on *A. curassavica* (Fig. 1), it was not possible to detect any cardenolide signals in the ectoperitrophic space, gut epithelium and hemolymph (see magnified view of the highlighted ROI in Figs. 3 S16 and S18), suggesting rapid clearance of cardenolides distributed in the caterpillars’ body fluids. For *E. core*, we did not detect any cardenolides after purging the gut lumen overnight with the cardenolide-free plant *O. coeruleum* (Fig. S18).

### Selectivity, modification and storage of cardenolides in *D. plexippus*

We studied the selectivity of sequestration as well as the transport and storage of sequestered cardenolides by MSI experiments with a higher spatial resolution (5 μm step size) to improve resolution on the tissue level. Despite being minor components in *A. curassavica* leaves (Fig. S19 for LC-MS-based quantification), calotropin/calactin (Fig. 4b), frugoside (Fig. S20b), and calotoxin (Fig. S20c) were the most abundant cardenolides in the gut lumen and integument of the monarch caterpillar. In contrast, although being one of the most abundant cardenolides in *A. curassavica* (LC-MS based quantification revealed 5.8 x higher concentration relative to calotropin, n = 3; Fig. S19), asclepin was exclusively detected at the fringes of the leaf particles in the gut lumen and was not sequestered into the body tissues (Fig. 4c).

**Figure 4.**
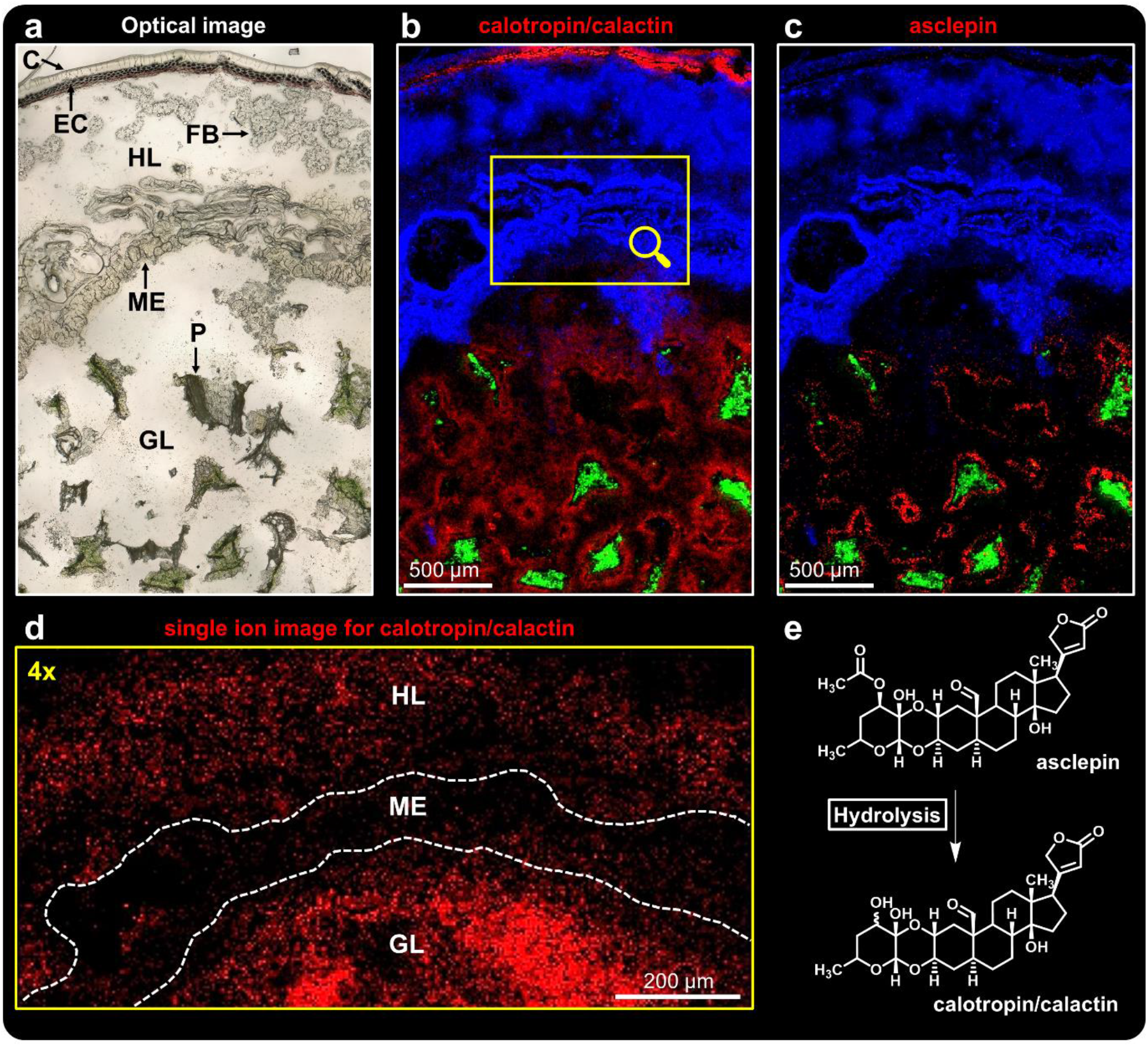
High-resolution AP-SMALDI MSI (5 μm step size) of a transversal *D. plexippus* (biological replicate 7) section. (**a**) Optical image of the analyzed region of interest. C: cuticle, EC: epidermal cells, HL: hemolymph, FB: fat body, ME: midgut epithelium, GL: gut lumen, P: *A. curassavica* plant material. (**b**,**c**) RGB overlay images showing the spatial distribution of (**b**) calactin/calotropin ([M+K]^+^, red) at *m/z* 571.2304, (**c**) asclepin ([M+K]^+^, red) at *m/z* 613.2427, and (**b,c**) pheophytin a ([M+K]^+^, green) at *m/z* 909.5290 and PE(36:3) ([M+K]^+^, blue) at *m/z* 780.4940. (**d**) Magnified view (4x) of the region outlined in **b**, showing the single-ion image for calactin/calotropin ([M+K]^+^) in red. Due to the high ion intensity in the integument, the intensity scale for this image was adjusted to visualize cardenolides in the midgut epithelium. (**e**) Putative degradation pathway of asclepin in *D. plexippus* gut.

Sequestered cardenolides were primarily stored in the epithelial cells of the integument (Figs. 4 and S20) and not in the cuticle (see ref. 51 for a histological section of monarch integument). While uscharidin and voruscharin were the dominant cardenolides in *A. curassavica*, their concentration in the integument was 5.2 x and 50 x lower compared to calotropin (n = 3; Fig. S19). However, although only at low amounts, we were able to visualize both toxins in the monarch caterpillar integument (Fig. S20e,f). Interestingly while not showing signals in the gut lumen, calotropagenin, which was the cardenolide with the lowest concentration in *A. curassavica* leaves (Fig. S19), was detected in the *D. plexippus* integument (Fig. S27a). Despite being an abundant compound in *A. curassavica* (Fig. S19) like asclepin, the biglucoside uzarin was not sequestered by monarch caterpillars (Fig. S20f).

Sequestration of plant toxins is generally assumed to be mediated via the epithelium of the midgut (Beran and Petschenka, 2022). In line with this prediction, we were able to detect calotropin/calactin with low abundance in the midgut epithelium of our transversal sections (Fig. 4d). We also monitored the distribution of cardenolides at the interface gut lumen, midgut epithelium, and hemolymph based on longitudinal caterpillar sections to address the molecular transport process in a larger spatial environment (Fig. 5). Here, we were able to visualize the spatial distribution of the dominant sequestered cardenolides calactin/calotropin (Fig. 5b), frugoside, (Fig. S21b) and calotoxin (Fig. S21c) within the epithelial tissue of the monarch midgut. In agreement with the putative role of the midgut epithelium as the transport organ for sequestered toxins mediating selectivity, the cardenolides uzarin (Fig. 5c) and asclepin (Fig. S21d) which were absent from the body cavity (i.e. not sequestered), were also not detected within the layer of midgut cells although they were present in the gut lumen. The cardenolide composition of the midgut epithelium tissue was further validated by LC-MS/MS experiments (Table 1).

**Figure 5.**
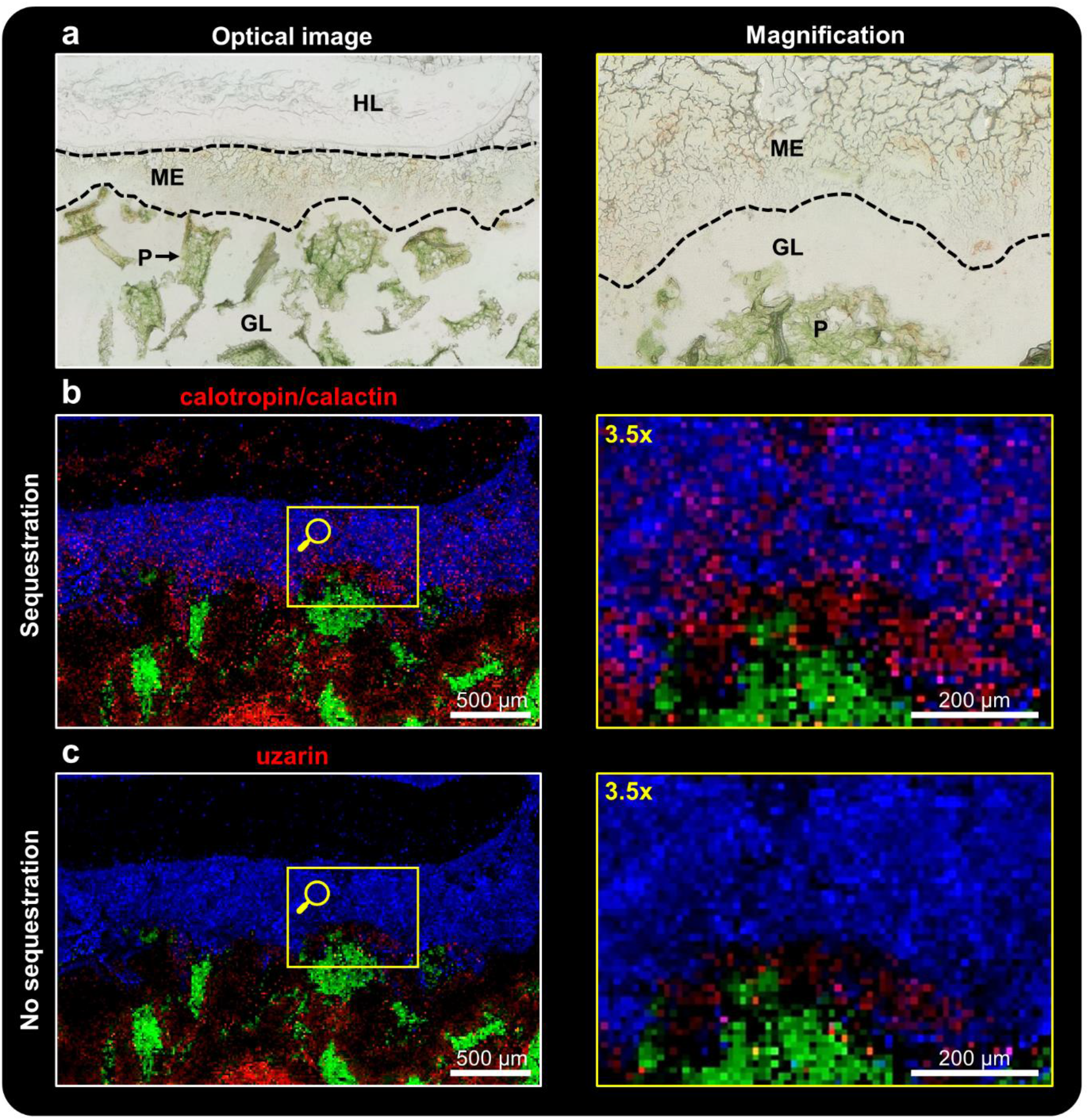
High-resolution AP-SMALDI MSI (10 μm step size) of a longitudinal *D. plexippus* (biological replicate 3) section. (**a**) Optical image of the analyzed region of interest and magnified view (3.5x) of the outlined region in **b**,**c**. P: *A. curassavica* plant material, GL: gut lumen, ME: midgut epithelium, HL: hemolymph. (**b**,**c**) RGB overlay images and magnified view of the outlined region showing the spatial distribution of calactin/calotropin ([M+K]^+^, red) at *m*/*z* 571.2304 in (**b**), uzarin ([M+K]^+^, red) at *m*/*z* 737.3149 in (**c**), pheophytin a ([M+K]^+^, green) at *m/z* 909.5290 and PE(36:3) ([M+K]^+^, blue) at *m/z* 780.4940.

**Table 1.** Overview of the 10 different cardenolides (Figure S8 for chemical structures) detected in *A. curassavica* leaves, gut content and various tissue types of *D. plexippus* and *E.core* (MG: midgut) using AP-SMALDI MSI and complementary LC-MS experiments. Note that native *A. curassavica* leaves were not analyzed via AP-SMALDI MSI.

| Compound | *A. curassavica*<br>leaf | *D. plexippus*<br>gut content | *E. core*<br>gut content | *D. plexippus*<br>MG epithelium | *D. plexippus*<br>integument | *E. core*<br>integument |
| --- | --- | --- | --- | --- | --- | --- |
| calotropagenin | +<sup>a</sup> | +<sup>a,b</sup> | +<sup>a</sup> | +<sup>a</sup> | +<sup>a,b</sup> | -<sup>a,b</sup> |
| uscharidin | +<sup>a</sup> | +<sup>a,b</sup> | +<sup>a,b</sup> | +<sup>a</sup> | +<sup>a,b</sup> | -<sup>a,b</sup> |
| calotropin | +<sup>a</sup> | +<sup>a,b</sup> | +<sup>a,b</sup> | +<sup>a,b</sup> | +<sup>a,b</sup> | -<sup>a,b</sup> |
| calactin | +<sup>a</sup> | +<sup>a,b</sup> | +<sup>a,b</sup> | +<sup>a,b</sup> | +<sup>a,b</sup> | -<sup>a,b</sup> |
| frugoside | +<sup>a</sup> | +<sup>a,b</sup> | +<sup>a,b</sup> | +<sup>a,b</sup> | +<sup>a,b</sup> | -<sup>a,b</sup> |
| calotoxin | +<sup>a</sup> | +<sup>a,b</sup> | +<sup>a,b</sup> | +<sup>a,b</sup> | +<sup>a,b</sup> | -<sup>a,b</sup> |
| asclepin | +<sup>a</sup> | +<sup>a,b</sup> | +<sup>a,b</sup> | -<sup>a,b</sup> | -<sup>a,b</sup> | -<sup>a,b</sup> |
| uscharin | +<sup>a</sup> | +<sup>a,b</sup> | +<sup>a</sup> | +<sup>a</sup> | +<sup>a,b</sup> | -<sup>a,b</sup> |
| voruscharin | +<sup>a</sup> | +<sup>a,b</sup> | +<sup>a</sup> | +<sup>a</sup> | +<sup>a,b</sup> | -<sup>a,b</sup> |
| uzarin | +<sup>a</sup> | +<sup>a,b</sup> | +<sup>a,b</sup> | -<sup>a,b</sup> | -<sup>a,b</sup> | -<sup>a,b</sup> |

+<sup>a</sup>: Detected via HPLC-MS
+<sup>a,b</sup>: Detected via HPLC-MS and AP-SMALDI MSI
-<sup>a,b</sup>: No detection via HPLC-MS and AP-SMALDI MSI

In addition, our high-resolution MSI analyses revealed that sequestered cardenolides appear in the monarch gut epithelium in a discrete, granular pattern, which was maintained after dispersion in the hemolymph (Figs. 5b, S21b,c). Notably, this pattern does not only apply to the distribution of cardenolides but also to other metabolites (Fig. S21, e.g. kaempferol-glucopyranoside, a ubiquitous plant secondary compound also occurring in milkweed).

We compared the distribution of cardenolides with the distribution of primary and other secondary metabolites, such as kaempferol-glucopyranoside (Fig. S21e), malvidin-glucoside (Fig. S21f), N-(1-deoxy-1-fructosyl)tyrosine (Fig. S21g), guanosine (Fig. S21h for *D. plexippus* and Fig. S22c for *E. core*) and unidentified disaccharides (Fig. S21i for *D. plexippus* and Fig. S22d for *E. core*) within the midgut epithelial tissue of *D. plexippus* and *E. core*. For both species, we observed spatial distributions similar to sequestered cardenolides in the monarch regarding the extraction from plant material and the transport across the gut epithelium. Therefore, utilizing the high spatial resolution ( ≤10 μm) provided by the AP-SMALDI MSI system, we were able to reveal the spatial organization of metabolite uptake at the sub-tissue level.

## Discussion

The demand for spatially-characterized biochemistry and molecular biology has grown rapidly over the recent years (Buchberger et al., 2018; Joo et al., 2008; Kherlopian et al., 2008; Spengler, 2015; van Hove et al., 2010). Traditional methods,such as immunofluorescence, require the labelling of biomolecules with fluorophores, which can be time-consuming, inefficient and is restricted to individual pre-known compounds (Yang et al., 2020). Moreover, labelling of molecules will most likely alter their physicochemical properties and therefore influence their tissue distribution. Despite its great potential, matrix-assisted laser desorption/ionization (MALDI) MSI was only rarely employed to study sequestration of plant toxins in insects (see Abdalsamee et al., 2014).

Our MSI-based spatial metabolomics approach allows for spatially-resolved, qualitative, and semi-quantitative analyses of metabolites and lipids in an untargeted fashion in their native state. We visualized diverging strategies of two closely-related milkweed butterfly species regarding the processing and uptake of plant toxins in the gut as well as the storage of sequestered compounds for defense. First, we demonstrated that the midgut lumen as the first physiological layer to mediate selectivity, plays a vital role concerning how *D. plexippus* and *E. core* cope with a toxic cardenolide diet. In contrast to the gut lumen of *D. plexippus* where cardenolides were found over the entire lumen, no extracted cardenolides were observed in between the leaf particles in the gut lumen of *E. core;* instead, cardenolides were exclusively detected at the fringes of ingested leaf material (Figs. 1, S1 and S2). This pattern most likely suggests immediate degradation of cardenolides extracted from plant material. We suggested degradation of cardenolides in *E. core* based on analyzed gut contents earlier (Petschenka and Agrawal, 2015) and the visualized lack of cardenolides in the liquid phase of the gut strongly supports our hypothesis and shows that degradation happens directly in the midgut lumen.

Whole-body MSI on longitudinal caterpillar sections and complementary HPLC-MS and HPLC-DAD experiments based on segments from freeze-dried caterpillar guts allowed for tracking cardenolides along the caterpillar gut passage. Specifically, we found that cardenolides accumulated in the midgut of *D. plexippus* while the concentration dropped in the hindgut, suggesting a hitherto undescribed mechanism of toxin partitioning in the monarch gut (Figs. 2, S8, S12 and S13). In contrast, for *E. core*, we detected constant cardenolide concentrations along the gut passage including the hind gut (Figs. 2, S12 and S13). These observations indicate that cardenolides are selectively retained in the monarch gut lumen resulting in higher local concentrations of preferentially-sequestered cardenolides, such as calotropin, calactin, frugoside. We speculate that the accumulation of cardenolides in the midgut creates a steep concentration gradient supporting efficient sequestration.

We conducted MSI experiments with caterpillars fed with the cardenolide-free diet *O*. *coeruleum* to purge their guts from milkweed cardenolides to further address the hypothesis of cardenolide retention in the monarch caterpillar gut. Consistently with our prediction, calotropin/calactin, frugoside and calotoxin were still abundant between *O*. *coeruleum* leaf particles in the gut lumen of monarch caterpillars, while in the gut lumen of *E. core*, no cardenolides were detected anymore after feeding on *O. coeruleum* leaves (Figs. 3, S16 and S18). We propose a mechanism analogous to adsorption chromatography, retaining cardenolides in the gut lumen while the food contents are passing by and are defecated. How this could be mediated mechanistically remains an open question and it is interesting to note, that the spatial distribution of cardenolide MSI signals seems to resemble the shape of the *O. coeruleum* leaf particles suggesting adhesion of cardenolides to *O. coeruleum* leaf particles in the gut lumen. Moreover, after purging there were no cardenolides detected anymore in the ectoperitrophic space which showed strong intensities in caterpillars actively feeding on *A. curassavica* (Fig. 1). This observation may indicate removal of cardenolides by sequestration via the midgut epithelium. Similarly, the body cavity was free of cardenolides suggesting rapid clearance one the supply from the gut lumen is halted.

Besides differences in the distribution, we also observed striking differences regarding the structural composition of cardenolides in the midgut of both caterpillar species. Remarkably, the stereoisomer ratio of calotropin/calactin in the *E. core* gut lumen (24:1) differed significantly from that found in the monarch gut lumen (1.2:1) and *A. curassavica* leaves (1.6:1; Figs. S14 and S15). Based on the inhibition of Na^+^/K^+^-ATPase, calactin was reported to be > 3x more toxic for *E. core* than its stereoisomer calotropin (Petschenka et al., 2018). Hence, we suggest that *E. core* prevents cardenolide intoxication by minimizing the concentration of calactin in the midgut which could be either mediated by converting calactin into a structurally different cardenolide, by degradation, or by prevention of calactin production from uscharidin contained in the ingested plant material (Seiber et al., 1980). Along the same lines, monarch caterpillars could maintain a high concentration of highly toxic calactin for defense.

Unlike calactin, calotropin, and frugoside, the predominant cardenolides in *A. curassavica* leaves, uscharidin, asclepin, voruscharin and uscharin, do not belong to the preferentially-sequestered cardenolides of the monarch, despite having high structural similarity to calotropin and calactin and sharing the same aglycon (calotropagenin) (Figs. 4, S19, S20). Surprisingly, asclepin, which only differs from calotropin/calactin by having an acetoxy-group (-OAc) instead of a hydroxy-group (-OH) in the sugar moiety, is not sequestered and instead, was exclusively detected at fringes of ingested *A. curassavica* leaf material (Fig. 4c). Given the high structural similarity to calotropin/calactin, it seems puzzling that the molecular mechanism underlying sequestration prohibits the uptake of asclepin. More likely, asclepin might be rapidly converted into calactin and/or calotropin instead of being not sequestered (Fig. 4e).

Similar observations regarding the metabolic conversion of structurally-related cardenolides into calotropin and calactin by the monarch caterpillar were already made decades ago for uscharidin (Marty and Krieger, 1984; Seiber et al., 1980) and recently, for voruscharin by Agrawal et al. (2021). Interestingly, both, uscharidin and voruscharin occured at similar concentrations in *A. curassavica* leaves like asclepin, (Fig. S19), but were very abundant in the monarch gut lumen (Fig. S20e,f). Consequently, we hypothesize that the conversion rate of asclepin into calactin and/or calotropin is significantly higher compared to uscharidin and voruscharin, suggesting a passive mechanism such as gut pH as observed for voruscharin (Agrawal et al., 2021) and not an involvement of enzymes as it was shown for uscharidin (Marty and Krieger, 1984).

The monarch Na^+^/K^+^-ATPase shows up to 94-fold higher resistance against cardenolides compared to non-adapted Na^+^/K^+^-ATPases (Petschenka et al., 2018). Remarkably, monarch Na^+^/K^+^-ATPase is only less than twofold more resistant to the thiazolidine-ring-containing cardenolides uscharin and voruscharin which dominate the cardenolide spectrum of *A. curassavica* leaves compared to porcine Na^+^/K^+^-ATPase (Agrawal et al., 2021). Consequently, the lack of sequestration of uscharin and voruscharin by monarch caterpillars, observed in this study, was interpreted as an adaptation to avoid toxicity. However, due to the high sensitivity of the employed AP-SMALDI MSI system, we were able to detect and visualize the accumulation of uscharin and voruscharin in the epidermal cells of the integument (Fig. S20e,f) and LC-MS-based quantification revealed 160 x (uscharin) and 50 x lower concentrations compared to the predominantly sequestered calotropin (Fig. S19). In conclusion, our data suggest that uscharin and voruscharin are sequestered but to a comparatively low extent. Alternatively to representing plant defense compounds specifically directed against the monarch butterfly, reduced sequestration and limited resistance of monarch Na^+^/K^+^-ATPase towards uscharin and voruscharin might be due to the rapid spontaneous degradation of voruscharin in the caterpillar gut which is also likely for uscharin due to the high structural similarity. Consequently, these compounds would not be available as a substrate for sequestration and not exert selection pressure on monarch Na^+^/K^+^-ATPase.

Our MSI analyses indicate that the midgut epithelium plays a critical role regarding the selectivity of sequestration by prohibiting the uptake of individual cardenolides such as uzarin (Fig. 5c). For uzarin, metabolism can be ruled out as a factor preventing sequestration, since this compound was found abundantly in the gut lumen and was observed in direct contact with the apical surface of the midgut epithelium (Fig. 5c). In other words, we suggest that the midgut epithelium discriminates against individual compounds by an unknown mechanism. The non-sequestered biglucoside uzarin (Fig. 5c) was the largest cardenolide detected and is comparatively polar, suggesting that size or polarity could be important determinants. Selective uptake of structurally different non-milkweed cardenolides was already demonstrated earlier (Frick and Wink, 1995). Here, we show that also milkweed cardenolides that actually occur in the diet of the monarch caterpillar are sequestered selectively.

Although sequestration of plant toxins was described for more than 275 insect species involving different classes of chemical compounds (Beran and Petschenka, 2022; Opitz and Müller, 2009), it is still largely unknown how plant toxins are transported across the insect gut epithelium (i.e. from the gut lumen into the body cavity). Using high-resolution MSI (10 μm step size), we were able to detect cardenolides within the midgut epithelium of monarch caterpillars. Notably, cardenolides appeared in a discrete granular pattern (Figs 5 and S21), suggesting that cardenolides are transported as aggregates and not as individual molecules. This pattern might indicate a vesicular transport via transcytosis, similar to what was described for the uptake of albumin across the midgut of the silkworm (*Bombyx mori*) (Casartelli et al., 2005). Since this granular pattern of cardenolides was already observed in the midgut lumen, the putative cardenolide vesicles may be plant derived. Although our histological sections were comparatively thick and likely contained several layers of cells complicating interpretation, the uniform distribution of the observed particles suggests that the transport occurs in a trans- and not in a paracellular fashion. It was suggested earlier that a carrier mechanism mediates the transport of cardenolides across the midgut epithelium in monarch caterpillars (Frick and Wink, 1995). How the cardenolide aggregations observed here could be aligned with a carrier-mediated process, however, remains an open question.

Remarkably, also other secondary plant metabolites such as polysaccharides and flavonoids as well as primary metabolites (e.g. guanosine) appeared in the same granular pattern within the midgut epithelium (Fig S21e-i), suggesting that vesicular uptake is a universal mode of uptake for various metabolites. A similar pattern was found for *E. core* regarding these metabolites but remarkably, cardenolides were not detected in the midgut epithelium (Fig. S22). Lipophilic and amphiphilic allelochemicals are expected to be sequestered into lipid aggregates (micelles) in the fluid phase of the gut of herbivorous insects proportionately to their lipophilicity (Barbehenn, 1999). Moreover, micelles formed by the aggregation of lysophospholipids, galactosyl glycerides, long chain fatty acids, and other amphiphilic and lipophilic compounds are known to represent the primary constituents of the non-aqueous phase of the midgut fluid in insect herbivores (Barbehenn, 1999). Therefore, it is likely that the granules which we observed represent micelles containing cardenolides and other metabolites. The observation of these putative micelles in the midgut epithelium of monarch caterpillars suggests that micelles composed of plant lipids and cardenolides which may cross the midgut epithelium by diffusion, represent the mode of transport by which cardenolides are sequestered.

It is unclear, however, how caterpillars of *E. core* discriminate against the uptake of cardenolides while apparently taking up other metabolites in a similar fashion. The peritrophic membrane of grasshoppers has been demonstrated to prevent the uptake of micelles containing various plant toxins including the cardenolide digitoxin by ultrafiltration (Barbehenn, 1999). This mechanism, however, is unlikely to explain non-sequestration of cardenolides in *E. core*, since cardenolide granules were observed in close contact with the midgut epithelium and thus were able to cross the peritrophic membrane surrounding the food bolus (Fig S22). It rather seems likely that degradation of cardenolides in the fluid-phase of the gut prevents sequestration of cardenolides in *E. core*.

Collectively, our spatially-resolved metabolomics approach revealed novel insight into the selectivity and the mechanism of cardenolide sequestration as well as of the location of sequestered cardenolides in monarch butterflies and demonstrates the potential of high-resolution AP-SMALDI MSI to explore insect-plant interaction biochemistry and to unravel the spatiotemporal fate of xenobiotics in insects (e.g. insecticides).

## Materials and Methods

### Chemicals, plants and insects

If not stated otherwise, all chemicals were purchased from Sigma-Aldrich (Steinheim, Germany). Caterpillars of *D. plexippus* (origin Portugal) and *E. core* (origin Southeast Asia) for AP-SMALDI MSI were obtained commercially, and colonies were maintained in cages in a greenhouse under ambient conditions. Caterpillars from both species were raised on potted *A. curassavica* plants (grown from seeds of commercial origin), grown in the same greenhouse and watered, fertilized and pruned as needed. For MSI experiments, actively-feeding final-instar larvae of *D. plexippus* and *E.core* were directly collected from the plants and stored at −80 °C for no longer than 2 weeks until sample preparation. For regional gut dissection experiments, caterpillars of *E. core* were raised in the greenhouse as well, while caterpillars of *D. plexippus* were raised on potted *A. curassavica* plants in a climate chamber (Fitotron^®^ SGC 120, Weiss Technik, Loughborough, UK) at 26°C (day) and 22°C (night) under a 16:8 h day/night cycle at 60% humidity. Caterpillars for these experiments were raised on individual plants and leaf samples from each plant were taken for cardenolide quantification.

For feeding experiments with cardenolide-free plants, monarch caterpillars (harvested from *A. curassavica* plants in the greenhouse) were fed with the cardenolide-free (Warashina and Shirota, 2021) *Oxypetalum coeruleum* (Apocynaceae) for three hours before sampling. In a preliminary trial, we fed flowers of *A. curassavica* (colored red and yellow) to a last-instar caterpillar of *D. plexippus* under ambient conditions. After one-hour, colored fecal pellets were observed indicating a full gut passage. Since caterpillar feeding activity on *O. coeruleum* was reduced compared to *A. curassavica*, three additional *D. plexippus* caterpillars of a similar size were allowed to feed on *O. coeruleum* for three hours under the same conditions. Caterpillars of *E. core* were quite hesitant to feed on *O. coeruleum* and only one of three caterpillars finally accepted *O. coeruleum* and was left on the leaves overnight. After the trials, caterpillars were frozen at −80°C and treated as described above. Although the time of *O. coeruleum* feeding was different for both caterpillar species, only leaf particles of *O. coeruleum* were visible in the histological sections indicating complete replacement of *A. curassavica* plant material by *O. coeruleum* tissue.

*D. plexippus* caterpillars for the LC-MS analyses of isolated caterpillar integuments were raised on potted *A. curassavica* plants in a greenhouse under ambient conditions and collected together while actively feeding.

### Sample preparation for MALDI mass spectrometry imaging

Before cryo-sectioning, caterpillars of *D. plexippus* and *E. core* were transferred to a cryomicrotome (HM525, Thermo Fisher Scientific, Walldorf, Germany) and allowed to warm up to −25 °C for 60 minutes. For transversal sectioning, each caterpillar was cut in half and fixed onto a metal target using freezing water as an adhesive medium (see Figure S23, for a schematic overview of the complete workflow). Subsequently, transversal sections with 40 μm thickness were obtained at −25 °C, thaw-mounted onto glass slides and stored at −80 °C until MSI analysis. All transversal sections were obtained between the first and the second pro-leg of the caterpillar, to ensure sectioning in the midgut region and to achieve consistent experimental conditions across all biological replicates of the two species.

Whole-insect sectioning (i.e. longitudinal sections) is challenging due to the large size of final instar *D. plexippus* and *E.core* caterpillars (3 to 5 cm in length). Thus, an MSI-compatible sample preparation protocol to obtain longitudinal sections of excellent quality was developed. First, a custom-made cryomold (24 mm × 50 mm × 30 mm) was filled to one-third of its volume with aqueous gelatin solution (8% (w/v) for *D. plexippus*, 6% (w/v) for *E. core*) and transferred to −25 °C for 20 minutes. Then, frozen caterpillars were placed onto the solidified gelatin block, covered with fresh gelatin solution and immediately snap-frozen in liquid nitrogen to prevent potential analyte delocalization and tissue damage induced by the hot gelatin (45 °C).

For sectioning, the frozen gelatin block was transferred to −25 °C for 60 minutes, subsequently taken out of the cryomold and fixed onto the metal target using freezing water as an adhesive medium as described above (see Figure S1). Longitudinal sections with 40 μm thickness were obtained at −25 °C for *D. plexippus* and −18 °C for *E. core*, thaw-mounted onto glass slides and subsequently stored at −80 °C until MSI analysis. Before matrix application, tissue sections were brought to room temperature in a desiccator for 30 minutes to avoid condensation of water at the sample surface. Optical images of tissue sections were obtained using a Keyence VHX-5000 digital microscope (Keyence Deutschland GmbH, Neu-Isenburg, Germany).

Among the MALDI matrices tested (positive-ion mode: 2,5-dihydroxybenzoic acid (2,5-DHB), α-cyano-4-hydroxycinnamic acid (CHCA), 2-mercaptobenzothiazol (MBT); negative-ion mode: 9-aminoacridine (9-AA), bis(dimethylamino)naphthalene (DMAN)), cardenolides were exclusively detected using 2,5-DHB in positive-ion mode. Before each MSI experiment, a matrix solution of 30 mg/mL 2,5-DHB (2,5-DHB, for synthesis, Merck, Darmstadt, Germany) in acetone/water at 1:1 (v/v) with 0.1% trifluoroacetic acid (TFA, for spectroscopy, AppliChem GmbH, Darmstadt, Germany) was freshly prepared. A volume of 100 μL DHB matrix solution for transversal sections, 130 μL for longitudinal sections and 70 μL for high-resolution MSI experiments (≤ 10 μm step size) was sprayed onto the sample surface at a flow rate of 10 μl/min (5 μl/min for high-resolution MSI experiments) using an ultrafine pneumatic sprayer system (SMALDIPrep, TransMIT GmbH, Giessen, Germany). The nebulizing nitrogen gas pressure was 1 bar and the rotation was set to 500 rpm. Crystal sizes (≤ 10 μm) and homogeneity of the matrix layer on the sample surface were controlled under the microscope before MSI analysis.

After MSI measurement, specific tissue sections were washed for 2 min with ethanol (100%) to remove the matrix layer and subsequently stained with hematoxylin and eosin stain (H&E) for histological classification (Fig. S24).

### Collection of regional gut samples, integuments, and sample preparation for HPLC analysis

Actively feeding last-instar caterpillars (= 5^th^ instar) of *D. plexippus* (n = 9) and *E. core* (n = 6) were collected from *A. curassavica* plants and immersed in crushed ice for 10 min. In addition, a mature leaf was collected from each plant (n = 9 for *D. plexippus* and n = 6 for *E. core*), frozen at −80°C and freeze-dried. After chilling, caterpillars were fixed in a dissection tray lined with Sylgard 184 (Dow Corning, Midland, MI, USA) using insect pins under ice-cold PBS (phosphate buffered saline; 154 mM NaCl, 5.6 Na_2_HPO_4_, 1.1 mM KH_2_PO4, pH 7.4; Roti®-CELL, Carl Roth, Karlsruhe, Germany). After decapitation, the integument was opened by a median cut along the dorsum of the caterpillar using micro-scissors. Next, the integument was fixed at both sides of the caterpillar and all tissues adhering to the gut or floating in the body cavity were removed (i.e. Malpighian tubules, salivary glands). Subsequently, the preparation was washed with ice-cold PBS by pouring fresh PBS into the dish at one side while removing the buffer with an automatic pipettor from the other side of the dish keeping the preparation permanently submersed under buffer. After washing, the preparation was frozen at −80°C and eventually freeze-dried for three days. Precipitated salts were carefully removed using a soft brush and the caterpillar gut including its contents was separated into foregut, four equally sized portions of midgut, and hindgut (see Fig. S25).

Gut portions were weighed using an analytical balance and extracted for HPLC-DAD and HPLC-ESI-MS analysis. Dry gut portions (i.e. the surrounding epithelium and the gut contents) were transferred into 2 mL screw-cap vials (Sarstedt AG & Co. KG, Nümbrecht, Germany) and 1 mL methanol containing 0.02 mg/mL of the internal standard digitoxin (Sigma-Aldrich, Taufkirchen, Germany) was added. After the addition of ca. 900 mg zirconia beads (ø 2.3 mm, BioSpec Products, Inc., Bartlesville, OK, USA), samples were homogenized in a Fast Prep™ homogenizer (MP Biomedicals, LLC, Solon, OH, US) for two 45-sec cycles at a speed of 6.5 m/sec. Subsequently, samples were centrifuged at 16,100 x g and supernatants were transferred into fresh vials. Extraction of the samples was repeated once as described above using 1 mL methanol without the internal standard and the second supernatant was combined with the first supernatant. After evaporating samples to dryness under a stream of N_2_, dry residues were dissolved in 200 μL methanol by agitation in the Fast-Prep-24 instrument (without the addition of beads). Before HPLC-DAD analysis, samples were filtered via Rotilabo®-syringe filters (nylon, 0.45 μm pore size, ø 13 mm, Carl Roth GmbH & Co. KG, Karlsruhe, Germany).

Leaf samples of *A. curassavica* from the experiment with *D. plexippus* were processed in the same fashion as described for the gut portions using a subset of roughly 50 mg of leaf material for extraction (49.4 – 52.8 mg). For leaf samples from the experiment with *E. core*, whole freeze-dried leaves were homogenized in 15 ml tubes containing two ceramic sphere beads (MP Biomedicals, Eschwege, Germany). We added 2 ml of methanol, containing 0.2 mg/ml of the internal standard digitoxin and samples were homogenized in the Fast-Prep-24 instrument as described above. After centrifugation (10 min at 1,000 x g) and removal of the supernatant, the extraction procedure was repeated with 2 ml pure methanol. Pooled supernatants were dried in a vacuum centrifuge and transferred into a 2-ml tube by washing the original tube with 3 x 500 μl methanol. Finally, the samples were processed as described above for HPLC analysis.

For the analyses of *D. plexippus* caterpillar integuments, three final-instar caterpillars were chilled on ice for 10 min. Subsequently, integuments were dissected, quickly rinsed in PBS, and adhering PBS was removed by pulling the samples over the edge of a glass beaker. Integument samples were stored at −80 °C and extracted as described above for gut portions but resuspended in 100 μl methanol instead.

### Instrumentation for MALDI mass spectrometry imaging

All MSI measurements were performed using an autofocusing AP-SMALDI5 AF ion source (TransMIT GmbH, Giessen, Germany) coupled to a Q Exactive HF Orbitrap mass spectrometer (Thermo Fisher Scientific, Bremen, Germany). For desorption/ionization, 50 laser pulses per pixel at 343 nm and a pulse rate of 100 Hz were focused perpendicular to the sample surface to an effective ablation spot diameter of ~5 μm (Fig. S26 for laser burn patterns of MSI experiments conducted with various step sizes). Whole-body and high-resolution (5 μm to 10 μm step size) MSI experiments were performed using the 3D-surface mode (“pixel-wise autofocusing”) to keep the MALDI laser focus, fluence and ablation spot size constant for varying sample surface characteristics (e.g. ingested plant material in the gut lumen or integument of the caterpillar, Fig S27) and to handle the sample tilt for the longitudinal caterpillar sections (~4 cm in length). The AP-SMALDI5 AF ion source was operated using the SMALDIControl software package (TransMIT GmbH, Giessen, Germany). The step size of the XYZ sample stage was set to the desired pixel size.

For all presented AP-SMALDI MSI experiments, the mass spectrometer was operated in positive-ion mode in a mass-to-charge-ratio (*m*/*z*) range of 250 to 1000 at a mass resolution of 240,000 at *m/z* 200. Internal lock-mass calibration was performed by using the ion signal from the DHB matrix cluster at *m/z* 716.12451 ([5DHB-4H_2_O+NH_4_]_+_), resulting in a mass accuracy of less than 2 ppm root mean square error (RMSE) for the whole image (Fig. S28). The scan speed for MSI experiments was about 1.4 pixel/s. The acceleration voltage was set to 3 kV. The ion injection time was set to 500 ms. The capillary temperature was 250 °C, and the S-lens level was set to 100 arbitrary units.

### Instrumentation for HPLC-DAD and HPLC-ESI-MS

All HPLC-ESI-MS experiments were performed using a Dionex UltiMate 3000 HPLC instrument (Thermo Fisher Scientific, Massachusetts, USA) coupled to a Q Exactive HF-X Orbitrap mass spectrometer (Thermo Fisher Scientific, Bremen, Germany). Analytes were separated on a Kinetex® C18 reversed-phase column (2.6 μm, 100 x 2.1 mm, Phenomenex, Torrance, USA). The injection volume was 15 μL, and the column compartment was set to 30 °C and 50 °C, respectively. Mobile phase A was water (0.1 % FA) and mobile phase B was acetonitrile (0.1 % FA). The following gradient was used at a flow rate of 0.5 mL/min: 0-2 min, 10% B; 2-20 min, 20-70% B; 20-25 min, 70-95% B; 25-30 min, 95% B; 30-35 min, 95-10% B. The mass spectrometer was operated in positive-ion mode in a mass-to-charge-ratio (*m*/*z*) range of 250 to 1000 at a mass resolution of 240,000 at *m/z* 200. The following HESI-source parameters were applied: spray voltage (+), 3.5 kV; capillary temperature, 300 °C; sheath gas flow rate, 35 psi; aux gas flowrate, 12 psi; aux gas heater temperature, 150 °C. Normalized collision energy (NCE) of 25% was used for fragmentation with z = 1 as default charge state. In total, three biological replicates of every sample type were measured (regional gut samples from *D. plexippus* and *E. core; D. plexippus* midgut tissue, integument; *A. curassavica* leaf).

For HPLC-DAD analyses, we injected 15 μl of extract into an Agilent 1100 series HPLC (Agilent Technologies, Santa Clara, USA) and compounds were separated on an EC 250/4.6 NUCLEODUR® C18 Gravity column (3 μm, 250 mm x 4.6 mm, Macherey-Nagel, Düren, Germany). Cardenolides were eluted at a constant flow of 0.7 ml/min at room temperature with an acetonitrile–H2O gradient as follows: 0–1 min 20% acetonitrile, 31 min 30% acetonitrile, 47 min 50% acetonitrile, 49 min 95% acetonitrile, 54 min 95% acetonitrile, 55 min 20% acetonitrile reconditioning for 10 min at 16% acetonitrile. UV-absorbance spectra were recorded from 200 to 400 nm with a diode array detector.

### Data processing

Xcalibur (Thermo Fisher Scientific, Massachusetts, USA) was used to display mass spectra. Ion images of selected *m/z* values were generated using MIRION imaging software (Paschke et al., 2013) (TransMIT GmbH, Giessen, Germany). The general principle for image generation is displayed in Figure S29. The ion-selection bin width (Δ(*m*/*z*)) of the images, generated from the MS data was set to *Δ(m/z) =* 0.01. Images were normalized to the base pixel (highest intensity of *m/z* bin) per image if not stated differently. No further data manipulation steps, such as smoothing or interpolation, were used. RGB MS images were obtained by selecting and overlaying three different images for the red-green-blue channels. The resulting images were adjusted in brightness for optimal visualization. MS imaging data were also converted to imzML using RAW2IMZML (TransMIT GmbH, Giessen, Germany), and MSiReader (Robichaud et al., 2013) was used to extract the ion intensities of specific *m*/*z* bins for defined regions of interest in the image. Metabolites and lipids were assigned and identified based on accurate mass measurements with a mass tolerance of less than 2 ppm RMSE for the whole image, MS/MS experiments and database matches (Palmer et al., 2017). For instance, calotropin/calactin ([M+K]^+^) at *m/z* 571.2304 was detected with a mass error of 0.5 ppm, and a root mean square error (RMSE) of 1.6 ppm (see Fig. S28a for RMSE plot).

For relative quantification of cardenolides based on MSI data, we defined seven regions of interest *in silico* (focused on *A. curassavica* leaf material and surrounding cardenolide signals; see Fig. S9 for a representative example based on biological replicate 1 of *D. plexippus* and *E. core*) per biological replicate and extracted the ion intensities for calotropin/calactin, calotoxin, uscharidin, frugoside and asclepin. Throughout all MSI experiments, pheophytin a was homogenously distributed and detected with marginal intensity variance in *A. curassavica* leaf pieces in *D. plexippus* and *E. core* gut lumen. Thus, for better comparison, we utilized the plant tissue marker pheophytin a as an internal standard and normalized the average cardenolide signal abundance to the average pheophytin a signal abundance for the respective region of interest.

For HPLC-MS based quantification, MZmine 2 (Pluskal et al., 2010) was used for preprocessing and extracting the area under the peak (AUP) of cardenolide features in the chromatogram. Subsequently, the respective cardenolide AUP was normalized to the AUP of the internal standard (Digitoxin) and the corresponding extracted tissue weight.

Data from HPLC-DAD analyses were evaluated using Agilent ChemStation (Rev. B.04.03, Agilent Technologies, Santa Clara, USA). Peaks with symmetrical absorption maxima between 216 and 222 nm (Malcolm and Zalucki, 1996) were interpreted as cardenolides and integrated at 218 nm. For gut samples obtained from dissections of freeze-dried caterpillars, we quantified cardenolides based on the peak area of the known concentration of the internal standard digitoxin. For leaf samples of *A. curassavica*, digitoxin was co-eluting with an endogenous cardenolide peak. Therefore, we used the mean of all digitoxin peak areas from the *E. core* gut samples (n = 36) which were run in the same batch for calibration of the leaf samples. Leaf samples collected during our experiment with *D. plexippus* were measured several months after the *D. plexippus* gut samples. Hence, cardenolides in these leaf samples were quantified using a digitoxin calibration curve which was measured shortly after the leaf samples were analyzed. Cardenolide concentrations of all leaf samples were corrected for the known amount of digitoxin which had been added initially as an internal standard. For caterpillar gut samples and leaf samples of *A. curassavica*, only peaks were considered which were present and had a clear cardenolide spectrum in at least 50 % of all samples. The observed pattern of results (see results section) did not change, when all peaks showing a clear cardenolide spectrum in each sample were considered for quantification.

### Statistical Analysis

We compared the cardenolide concentrations of dissected gut portions and leaf material using the standard least squares model in JMP Pro 15 (SAS Institute, Cary, NC, USA) including caterpillar individual as a model effect. For the *D. plexippus* dataset, data were log10-transformed to achieve normality of residuals and homogeneity of variance. For comparing intensities of cardenolide signals *in silico*, we calculated means of all seven ROIs for each cardenolide in each caterpillar. Means for each cardenolide were compared between caterpillar species using t-tests assuming unequal variances in JMP Pro 15. P-values < 0.05 were considered statistically significant.

## Supporting information

Dreisbach_et_al_supplementary_information

## Acknowledgements

We thank Hermann Falkenhahn, Johanna Weber, and Sabrina Stiehler for insect rearing, growing of plants and collecting samples. Technical support by Thermo Fisher Scientific GmbH (Bremen, Germany) and TransMIT GmbH (Giessen, Germany) are gratefully acknowledged. This research was funded by DFG grant PE 2059/3-1 to G.P. and the LOEWE Program of the State of Hesse by funding the LOEWE Center for Insect Biotechnology and Bioresources.

## Author contributions

G.P., B.S. and D.D. designed research, D.D., G.P., A.B., L.T. and D.B. performed research, D.D. and G.P. analyzed data, D.D., G.P., B.S., and A.V. wrote the paper.

## Data availability

All underlying data will be made available upon acceptance of the manuscript.

## References

Abdalsamee MK, Giampà M, Niehaus K, Müller C. 2014. Rapid incorporation of glucosinolates as a strategy used by a herbivore to prevent activation by myrosinases. Insect Biochem Mol Biol 52:115–123. doi:https://doi.org/10.1016/j.ibmb.2014.07.002

Agrawal AA, Böröczky K, Haribal M, Hastings AP, White RA, Jiang R-W, Duplais C. 2021. Cardenolides, toxicity, and the costs of sequestration in the coevolutionary interaction between monarchs and milkweeds. Proc Natl Acad Sci 118:e2024463118. doi:10.1073/pnas.2024463118

Agrawal AA, Petschenka G, Bingham RA, Weber MG, Rasmann S. 2012. Toxic cardenolides: chemical ecology and coevolution of specialized plant–herbivore interactions. New Phytol 194:28–45. doi:https://doi.org/10.1111/j.1469-8137.2011.04049.x

Ayres MP, Clausen TP, MacLean Jr SF, Redman AM, Reichardt PB. 1997. Diversity of structure and antiherbivore activity in condensed tannins. Ecology 78:1696–1712.

Barbehenn R V. 1999. Non-absorption of ingested lipophilic and amphiphilic allelochemicals by generalist grasshoppers: The role of extractive ultrafiltration by the peritrophic envelope. Arch Insect Biochem Physiol 42:130–137. doi:https://doi.org/10.1002/(SICI)1520-6327(199910)42:2<130::AID-ARCH3>3.0.CO;2-C

Beran F, Pauchet Y, Kunert G, Reichelt M, Wielsch N, Vogel H, Reinecke A, Svatoš A, Mewis I, Schmid D, Ramasamy S, Ulrichs C, Hansson BS, Gershenzon J, Heckel DG. 2014. &lt;em&gt;Phyllotreta striolata&lt;/em&gt; flea beetles use host plant defense compounds to create their own glucosinolate-myrosinase system. Proc Natl Acad Sci 111:7349 LP – 7354. doi:10.1073/pnas.1321781111

Beran F, Petschenka G. 2022. Sequestration of Plant Defense Compounds by Insects: From Mechanisms to Insect–Plant Coevolution. Annu Rev Entomol 67:163–180. doi:10.1146/annurev-ento-062821-062319

Brower LP, Ryerson WN, Coppinger LL, Glazier SC. 1968. Ecological chemistry and the palatability spectrum. Science (80-) 161:1349–1350.

Buchberger AR, DeLaney K, Johnson J, Li L. 2018. Mass Spectrometry Imaging: A Review of Emerging Advancements and Future Insights. Anal Chem 90:240–265. doi:10.1021/acs.analchem.7b04733

Casartelli M, Corti P, Giovanna Leonardi M, Fiandra L, Burlini N, Pennacchio F, Giordana B. 2005. Absorption of albumin by the midgut of a lepidopteran larva. J Insect Physiol 51:933–940. doi:https://doi.org/10.1016/j.jinsphys.2005.04.008

Cresswell JE, Merritt SZ, Martin MM. 1992. The effect of dietary nicotine on the allocation of assimilated food to energy metabolism and growth in fourth-instar larvae of the southern armyworm, Spodoptera eridania (Lepidoptera: Noctuidae). Oecologia 89:449–453.

Dobler S, Dalla S, Wagschal V, Agrawal AA. 2012. Community-wide convergent evolution in insect adaptation to toxic cardenolides by substitutions in the Na, K-ATPase. Proc Natl Acad Sci 109:13040–13045.

Dobler S, Petschenka G, Wagschal V, Flacht L. 2015. Convergent adaptive evolution – how insects master the challenge of cardiac glycoside-containing host plants. Entomol Exp Appl 157:30–39. doi:https://doi.org/10.1111/eea.12340

Duffey SS. 1980. Sequestration of plant natural products by insects. Annu Rev Entomol 25:447–477.

Dussourd DE, Hoyle AM. 2000. Poisoned plusiines: toxicity of milkweed latex and cardenolides to some generalist caterpillars. Chemoecology 10:11–16. doi:10.1007/PL00001810

Fraenkel GS. 1959. The Raison d’Etre of Secondary Plant Substances: These odd chemicals arose as a means of protecting plants from insects and now guide insects to food. Science (80-) 129:1466–1470.

Frick C, Wink M. 1995. Uptake and sequestration of ouabain and other cardiac glycosides in Danaus plexippus (Lepidoptera: Danaidae): evidence for a carrier-mediated process. J Chem Ecol 21:557–575.

Geier B, Oetjen J, Ruthensteiner B, Polikarpov M, Gruber-Vodicka H, Liebeke M. 2021. Connecting structure and function from organisms to molecules in small animal symbioses through chemo-histo-tomography. bioRxiv 2009–2020.

Geier B, Sogin EM, Michellod D, Janda M, Kompauer M, Spengler B, Dubilier N, Liebeke M. 2020. Spatial metabolomics of in situ host–microbe interactions at the micrometre scale. Nat Microbiol 5:498–510.

Holzinger F, Wink M. 1996. Mediation of cardiac glycoside insensitivity in the monarch butterfly (Danaus plexippus): role of an amino acid substitution in the ouabain binding site of Na+, K+-ATPase. J Chem Ecol 22:1921–1937.

Iwama T, Kano K, Saigusa D, Ekroos K, van Echten-Deckert G, Vogt J, Aoki J. 2021. Development of an On-Tissue Derivatization Method for MALDI Mass Spectrometry Imaging of Bioactive Lipids Containing Phosphate Monoester Using Phos-tag. Anal Chem 93:3867–3875.

Joo C, Balci H, Ishitsuka Y, Buranachai C, Ha T. 2008. Advances in Single-Molecule Fluorescence Methods for Molecular Biology. Annu Rev Biochem 77:51–76. doi:10.1146/annurev.biochem.77.070606.101543

Kadesch P, Hollubarsch T, Gerbig S, Schneider L, Silva LMR, Hermosilla C, Taubert A, Spengler B. 2020a. Intracellular parasites Toxoplasma gondii and Besnoitia besnoiti, Unveiled in single host cells using AP-SMALDI MS imaging. J Am Soc Mass Spectrom 31:1815–1824.

Kadesch P, Quack T, Gerbig S, Grevelding CG, Spengler B. 2020b. Tissue-and sex-specific lipidomic analysis of Schistosoma mansoni using high-resolution atmospheric pressure scanning microprobe matrix-assisted laser desorption/ionization mass spectrometry imaging. PLoS Negl Trop Dis 14:e0008145.

Kadesch P, Quack T, Gerbig S, Grevelding CG, Spengler B. 2019. Lipid Topography in Schistosoma mansoni Cryosections, Revealed by Microembedding and High-Resolution Atmospheric-Pressure Matrix-Assisted Laser Desorption/Ionization (MALDI) Mass Spectrometry Imaging. Anal Chem 91:4520–4528.

Karageorgi M, Groen SC, Sumbul F, Pelaez JN, Verster KI, Aguilar JM, Hastings AP, Bernstein SL, Matsunaga T, Astourian M, Guerra G, Rico F, Dobler S, Agrawal AA, Whiteman NK. 2019. Genome editing retraces the evolution of toxin resistance in the monarch butterfly. Nature 574:409–412. doi:10.1038/s41586-019-1610-8

Kazana E, Pope TW, Tibbles L, Bridges M, Pickett JA, Bones AM, Powell G, Rossiter JT. 2007. The cabbage aphid: a walking mustard oil bomb. Proc R Soc B Biol Sci 274:2271–2277. doi:10.1098/rspb.2007.0237

Kherlopian AR, Song T, Duan Q, Neimark MA, Po MJ, Gohagan JK, Laine AF. 2008. A review of imaging techniques for systems biology. BMC Syst Biol 2:1–18.

Koestler M, Kirsch D, Hester A, Leisner A, Guenther S, Spengler B. 2008. A high-resolution scanning microprobe matrix-assisted laser desorption/ionization ion source for imaging analysis on an ion trap/Fourier transform ion cyclotron resonance mass spectrometer. Rapid Commun Mass Spectrom An Int J Devoted to Rapid Dissem Up-to-the-Minute Res Mass Spectrom 22:3275–3285.

Koiwa H, Bressan RA, Hasegawa PM. 1997. Regulation of protease inhibitors and plant defense. Trends Plant Sci 2:379–384.

Kompauer M, Heiles S, Spengler B. 2017a. Atmospheric pressure MALDI mass spectrometry imaging of tissues and cells at 1.4-μm lateral resolution. Nat Methods 14:90–96. doi:10.1038/nmeth.4071

Kompauer M, Heiles S, Spengler B. 2017b. Autofocusing MALDI mass spectrometry imaging of tissue sections and 3D chemical topography of nonflat surfaces. Nat Methods 14:1156–1158.

Malcolm SB, Zalucki MP. 1996. Milkweed latex and cardenolide induction may resolve the lethal plant defence paradoxProceedings of the 9th International Symposium on Insect-Plant Relationships. Springer. pp. 193–196.

Malcom S, Rothschild M. 1983. A danaid mullerian mimic, Euploea core amymone (Cramer) lacking cardenolides in the pupal and adult stages. Biol J Linn Soc 19:27–33. doi:10.1111/j.1095-8312.1983.tb00774.x

Marty MA, Krieger RI. 1984. Metabolism of uscharidin, a milkweed cardenolide, by tissue homogenates of monarch butterfly larvae, Danaus plexippus L. J Chem Ecol 10:945–956.

Narberhaus I, Zintgraf V, Dobler S. 2005. Pyrrolizidine alkaloids on three trophic levels-evidence for toxic and deterrent effects on phytophages and predators. Chemoecology 15:121–125.

Opitz SEW, Müller C. 2009. Plant chemistry and insect sequestration. Chemoecology 19:117–154.

Palmer A, Phapale P, Chernyavsky I, Lavigne R, Fay D, Tarasov A, Kovalev V, Fuchser J, Nikolenko S, Pineau C. 2017. FDR-controlled metabolite annotation for high-resolution imaging mass spectrometry. Nat Methods 14:57–60.

Parsons JA. 1965. A digitalis-like toxin in the monarch butterfly, Danaus plexippus L. J Physiol 178:290–304. doi:https://doi.org/10.1113/jphysiol.1965.sp007628

Paschke C, Leisner A, Hester A, Maass K, Guenther S, Bouschen W, Spengler B. 2013. Mirion—a software package for automatic processing of mass spectrometric images. J Am Soc Mass Spectrom 24:1296–1306.

Petschenka G, Agrawal AA. 2015. Milkweed butterfly resistance to plant toxins is linked to sequestration, not coping with a toxic diet. Proc R Soc B Biol Sci 282:20151865.

Petschenka G, Fei CS, Araya JJ, Schröder S, Timmermann BN, Agrawal AA. 2018. Relative selectivity of plant cardenolides for Na+/K+-ATPases from the monarch butterfly and non-resistant insects. Front Plant Sci 9:1424.

Pluskal T, Castillo S, Villar-Briones A, Orešič M. 2010. MZmine 2: Modular framework for processing, visualizing, and analyzing mass spectrometry-based molecular profile data. BMC Bioinformatics 11:395. doi:10.1186/1471-2105-11-395

Reichstein T von, Von Euw J, Parsons JA, Rothschild M. 1968. Heart poisons in the monarch butterfly. Science (80-) 161:861–866.

Robert CAM, Zhang X, Machado RAR, Schirmer S, Lori M, Mateo P, Erb M, Gershenzon J. 2017. Sequestration and activation of plant toxins protect the western corn rootworm from enemies at multiple trophic levels. Elife 6:e29307. doi:10.7554/eLife.29307

Robichaud G, Garrard KP, Barry JA, Muddiman DC. 2013. MSiReader: an open-source interface to view and analyze high resolving power MS imaging files on Matlab platform. J Am Soc Mass Spectrom 24:718–721.

Schatzmann HJ. 1953. Herzglykoside als Hemmstoffe fur den activen Kalium und Natrium-Transport durch die Erythrocytemembran. Helv physiol pharmacol Acta 11:346–354.

Seiber JN, Tuskes PM, Brower LP, Nelson CJ. 1980. Pharmacodynamics of some individual milkweed cardenolides fed to larvae of the monarch butterfly (Danaus plexippus L.). J Chem Ecol 6:321–339.

Spengler B. 2015. Mass Spectrometry Imaging of Biomolecular Information. Anal Chem 87:64–82. doi:10.1021/ac504543v

Spengler B, Hubert M. 2002. Scanning microprobe matrix-assisted laser desorption ionization (SMALDI) mass spectrometry: instrumentation for sub-micrometer resolved LDI and MALDI surface analysis. J Am Soc Mass Spectrom 13:735–748.

Spengler B, Hubert M, Kaufmann R. 1994. MALDI ion imaging and biological ion imaging with a new scanning UV-laser microprobeProceedings of the 42nd Annual Conference on Mass Spectrometry and Allied Topics. Chicago. p. 1041.

van Hove ERA, Smith DF, Heeren RMA. 2010. A concise review of mass spectrometry imaging. J Chromatogr A 1217:3946–3954.

Vaughan GL, Jungreis AM. 1977. Insensitivity of lepidopteran tissues to ouabain: physiological mechanisms for protection from cardiac glycosides. J Insect Physiol 23:585–589.

Warashina T, Shirota O. 2021. Tetracyclic Triterpenoids, Steroids and Lignanes from the Aerial Parts of *Oxypetalum caeruleum*. Chem Pharm Bull 69:226–231. doi:10.1248/cpb.c20-00789

Yang F, Chen J, Ruan Q, Saqib HSA, He W, You M. 2020. Mass spectrometry imaging: An emerging technology for the analysis of metabolites in insects. Arch Insect Biochem Physiol 103:e21643.

Yang Z-L, Nour-Eldin HH, Hänniger S, Reichelt M, Crocoll C, Seitz F, Vogel H, Beran F. 2021. Sugar transporters enable a leaf beetle to accumulate plant defense compounds. Nat Commun 12:2658. doi:10.1038/s41467-021-22982-8

Zhang R-R, Tian H-Y, Tan Y-F, Chung T-Y, Sun X-H, Xia X, Ye W-C, Middleton DA, Fedosova N, Esmann M, Tzen JTC, Jiang R-W. 2014. Structures, chemotaxonomic significance, cytotoxic and Na+,K+-ATPase inhibitory activities of new cardenolides from Asclepias curassavica. Org Biomol Chem 12:8919–8929. doi:10.1039/C4OB01545B

Züst T, Petschenka G, Hastings AP, Agrawal AA. 2019. Toxicity of Milkweed Leaves and Latex: Chromatographic Quantification Versus Biological Activity of Cardenolides in 16 Asclepias Species. J Chem Ecol 45:50–60. doi:10.1007/s10886-018-1040-3

