## Supplementary material for "Spatial metabolomics reveal divergent cardenolide processing in the monarch butterfly (*Danaus plexippus*) and the common crow (*Euploea core*)": Dreisbach_et_al_supplementary_information

**Supplementary Table 1**. Overview of the biological replicates of *D. plexippus* and *E. core*, which were analysed via AP-SMALDI MSI. Plant diet, orientation of cryo-sectioning and the figures for the corresponding MSI results are mentioned.

| Biological replicate | Diet | Cryo-sectioning | MSI results |
|---|---|---|---|
| ***Danaus plexippus*** **1** | *A. curassavica* | transversal | Figs. 1, S2 |
| ***Danaus plexippus*** **2** | *A. curassavica* | transversal | Fig. S1 |
| ***Danaus plexippus*** **3** | *A. curassavica* | longitudinal | Figs. 2, 5 |
| ***Danaus plexippus*** **4** | *A. curassavica* | longitudinal | Fig. S8 |
| ***Danaus plexippus*** **5** | *A. curassavica,* purged with *O. coeruleum* | transversal | Figs. 3, S16 |
| ***Danaus plexippus*** **6** | *A. curassavica,* purged with *O. coeruleum* | transversal | Fig. S18 |
| ***Danaus plexippus*** **7** | *A. curassavica* | transversal | Figs. 4, S20 |
| ***Euploea core 1*** | *A. curassavica* | transversal | Figs. 1, S2 |
| ***Euploea core 2*** | *A. curassavica* | transversal | Fig. S1 |
| ***Euploea core 3*** | *A. curassavica* | longitudinal | Fig. 2 |
| ***Euploea core 4*** | *A. curassavica* | longitudinal | Fig. S8 |
| ***Euploea core 5*** | *A. curassavica,* purged with *O. coeruleum* | transversal | Fig. S18 |

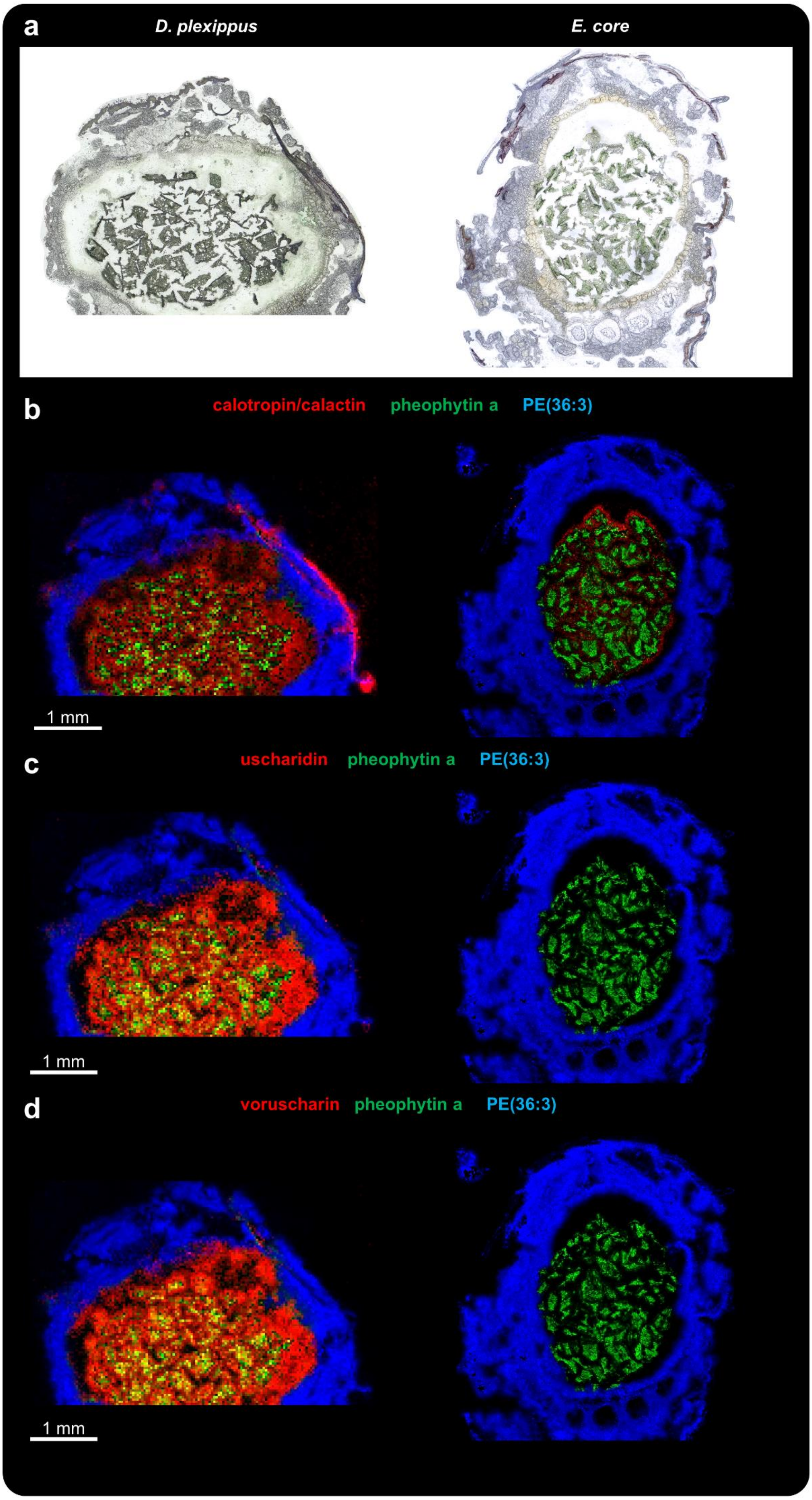
a
D. plexippus
E. core
b
calotropin/calactin
pheophytin a
PE(36:3)
1 mm
c
uscharidin
pheophytin a
PE(36:3)
1 mm
d
voruscharin
pheophytin a
PE(36:3)
1 mm

**Figure S1.** AP-SMALDI MSI of transversal sections of caterpillars of *D. plexippus* (biological replicate 2) and *E. core* (biological replicate 2). (**a**) Optical image of the transversal section of a fifth instar caterpillar of *D. plexippus* (left) and *E. core* (right) before matrix application. (**b-d**) Corresponding RGB overlay images obtained with 35 µm (*D. plexippus*) and 20 µm (*E. core*) step size showing the spatial distribution for the cardenolides (**b**) calotropin and/or its isomer calactin ($[M+K]^+$, red) at *m*/*z* 571.2304, (**c**) uscharidin ($[M+K]^+$, red) at *m*/*z* 569.2152, (**d**) voruscharin ($[M+K]^+$, red) at *m*/*z* 628.2346, (**b-d**) the chlorophyll derivative pheophytin a at *m*/*z* 909.5288 ($[M+K]^+$, green) as a chemical marker for plant tissue and the animal lipid PE(36:3) ($[M+K]^+$, blue) as a chemical marker for animal tissues. To demonstrate the differences regarding the cardenolide content between both species, the respective RGB ion images are normalized to the same intensity scale.

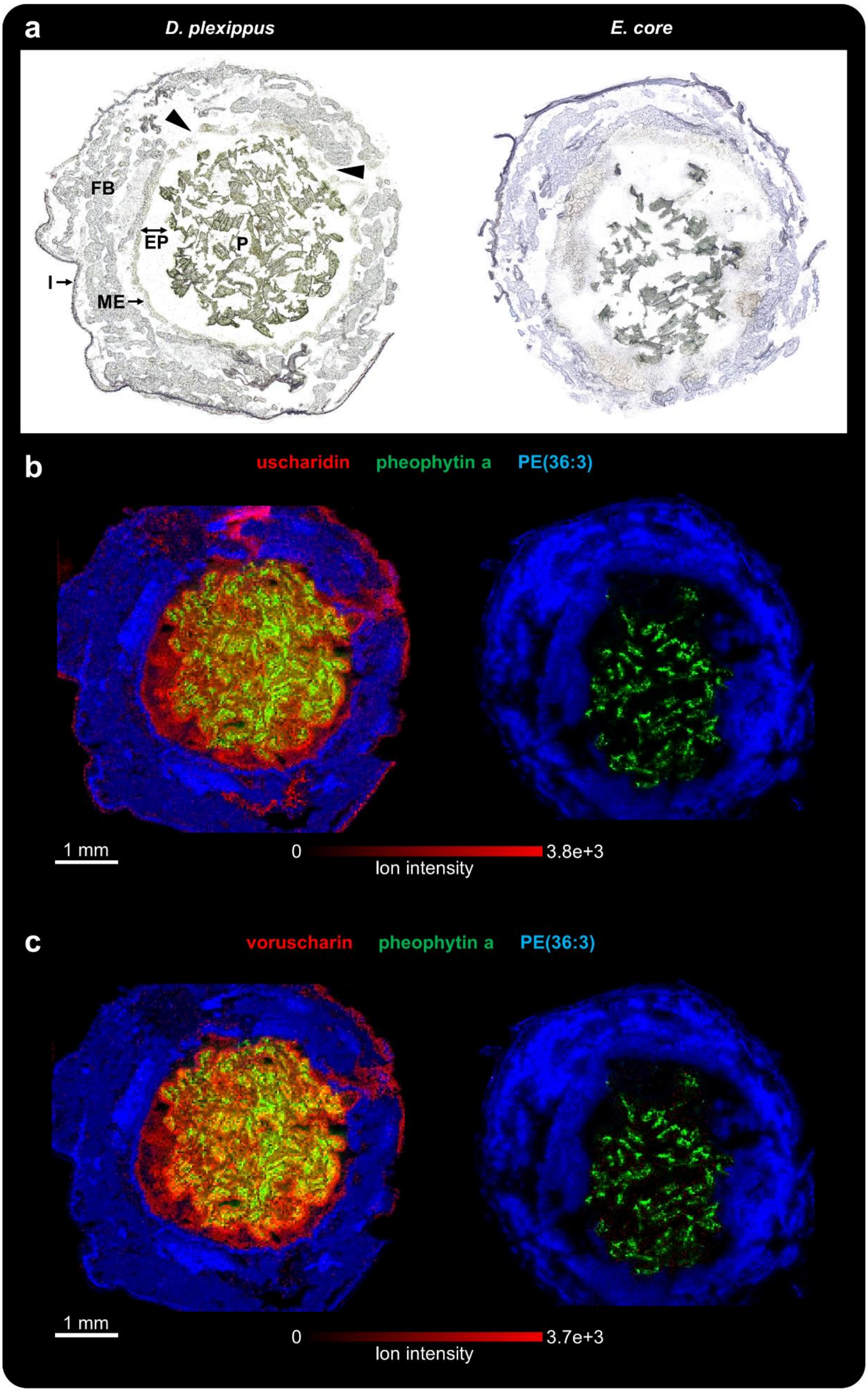


**Figure S2.** AP-SMALDI MSI of transversal sections of caterpillars of *D. plexippus* (biological replicate 1) and *E. core* (biological replicate 1). (**a**) Optical image of the transversal section of a fifth instar caterpillar of *D. plexippus* (left) and *E. core* (right) before matrix application. P:

plant material, EP: ectoperitrophic space, ME: midgut epithelium, FB: fat body, I: integument. (**b**) Corresponding RGB overlay images obtained with 35 µm (*D. plexippus*) and 20 µm (*E. core*) step size, showing the spatial distribution of the cardenolide (**b**) uscharidin ($[M+K]^+$, red) at *m*/*z* 569.2151, (**c**) voruscharin ($[M+K]^+$, red) at *m*/*z* 628.2346 and (**b,c**) the chlorophyll derivative pheophytin a at *m*/*z* 909.5288 ($[M+K]^+$, green) as a chemical marker for plant tissue and the animal lipid PE(36:3) ($[M+K]^+$, blue) as a chemical marker for animal tissues. To demonstrate the differences regarding the cardenolide content between both species, the respective RGB ion images are normalized to the same intensity scale. The *D. plexippus* gut epithelium was damaged at two areas (highlighted in the optical image), causing potential analyte delocalization in the corresponding hemolymph area.

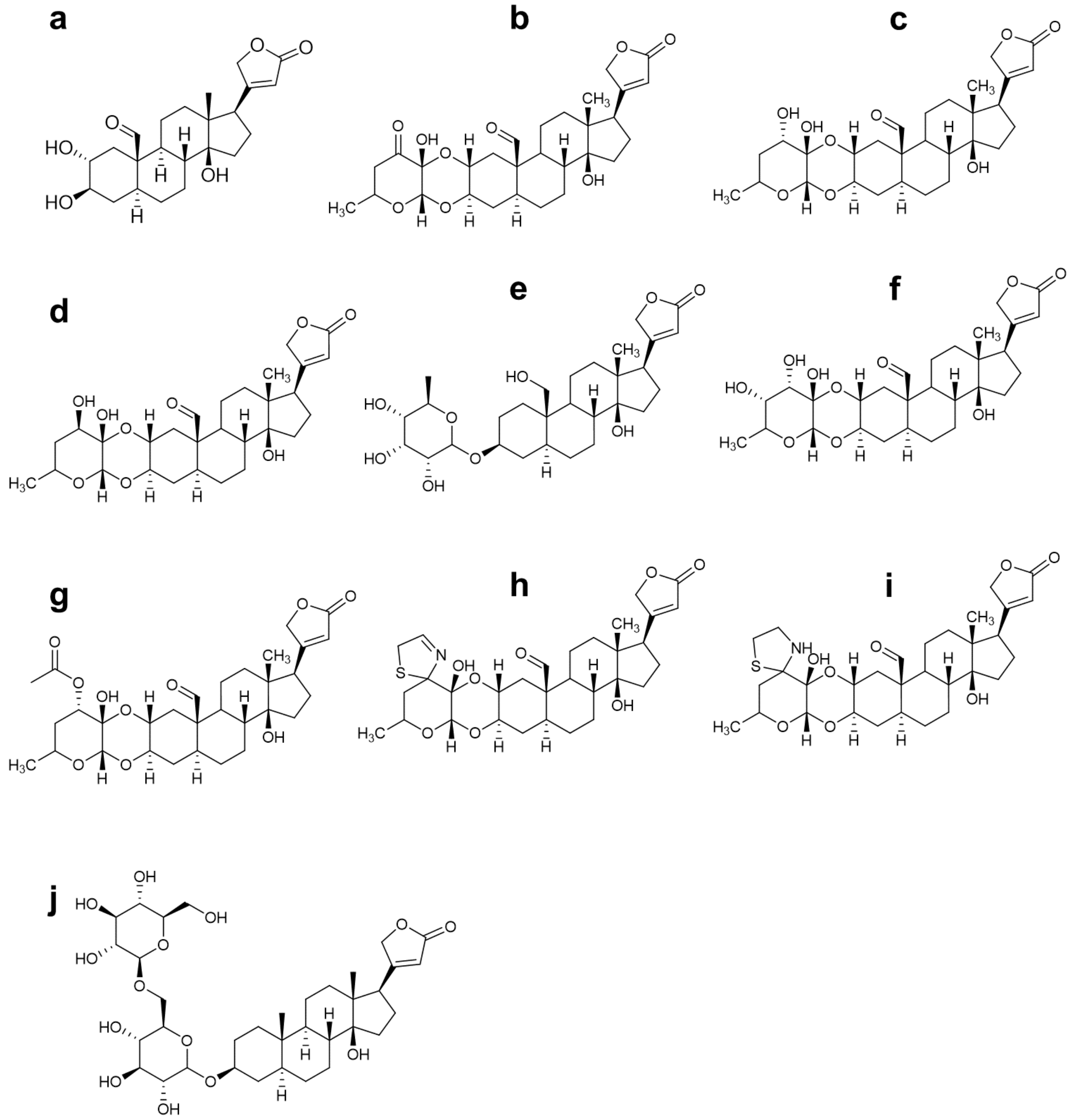


**Figure S3.** Chemical structures for all detected cardenolides. (**a**) calotropagenin, (**b**) uscharidin, (**c**) calotropin, (**d**) calactin, (**e**) frugoside, (**f**) calotoxin, (**g**) asclepin, (**h**) uscharin, (**i**) voruscharin, (**j**) uzarin.

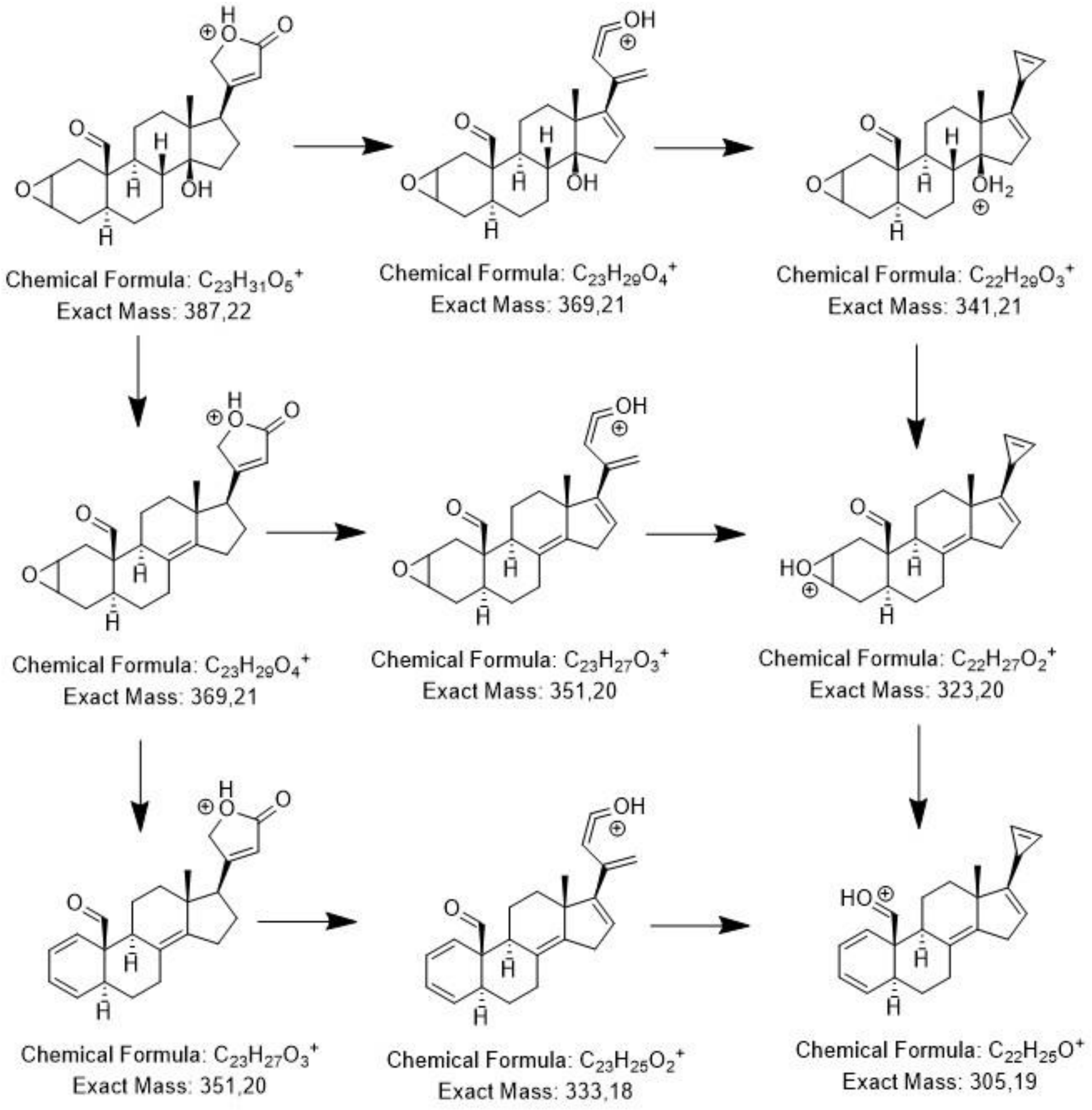


**Figure S4.** Proposed fragmentation pathway for cardenolides having an aldehyde function at C18 and a dioxane-linkage between the glycoside unit and the aglycone (i.e. all cardenolides detected with the exception of frugoside and uzarin). The fragmentation pathways including the chemical structures of the fragments are highlighted in the HPLC-MS/MS spectra of the respective cardenolide (Figs. S5-S7). For asclepin, uscharin and voruscharin, the characteristic fragments based on the glycoside unit are also displayed.

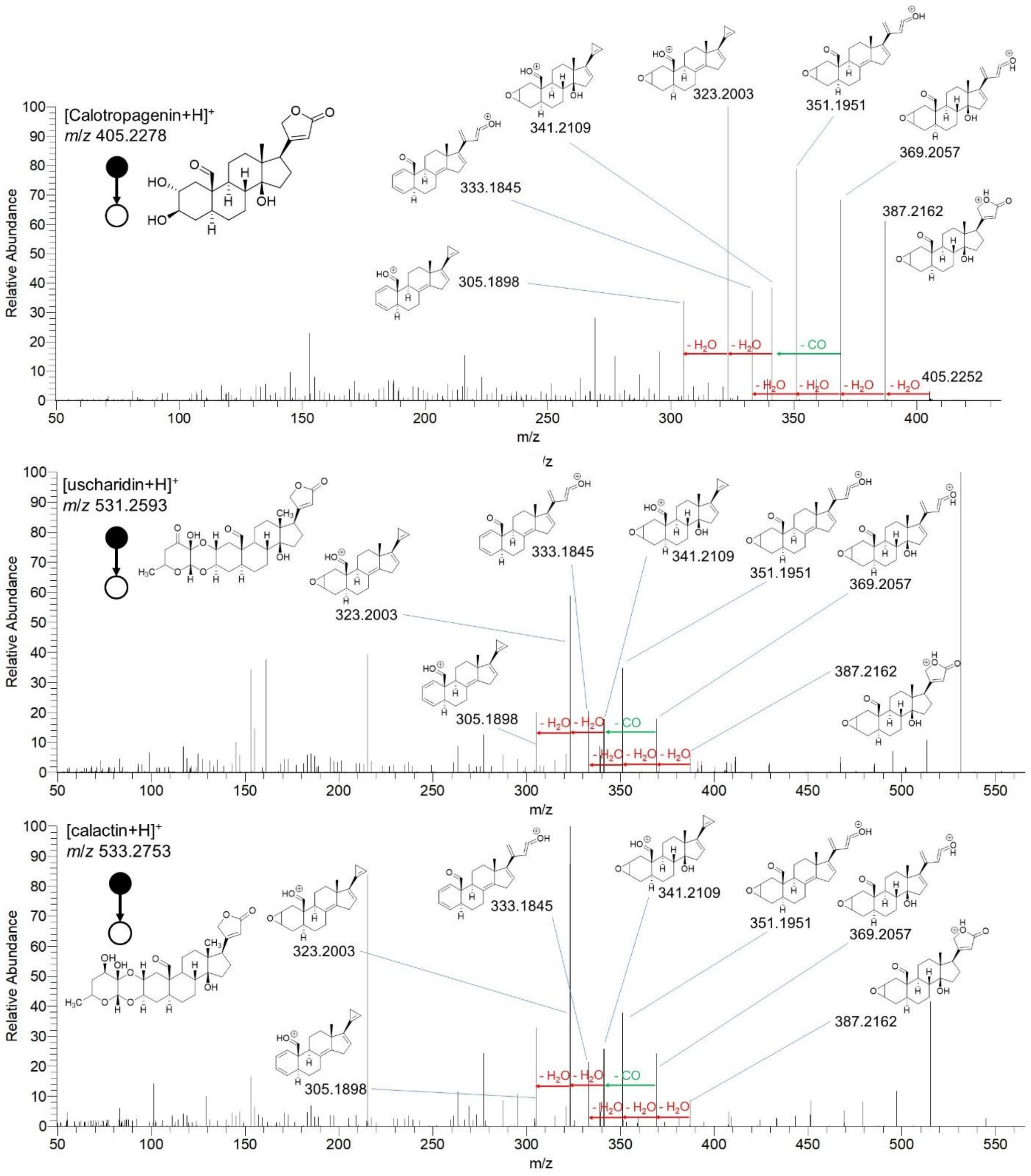


**Figure S5**. HPLC-MS/MS spectra for calotropagenin, uscharidin and calactin to confirm imaging annotations. Fragmentation pathways including characteristic fragments are highlighted.

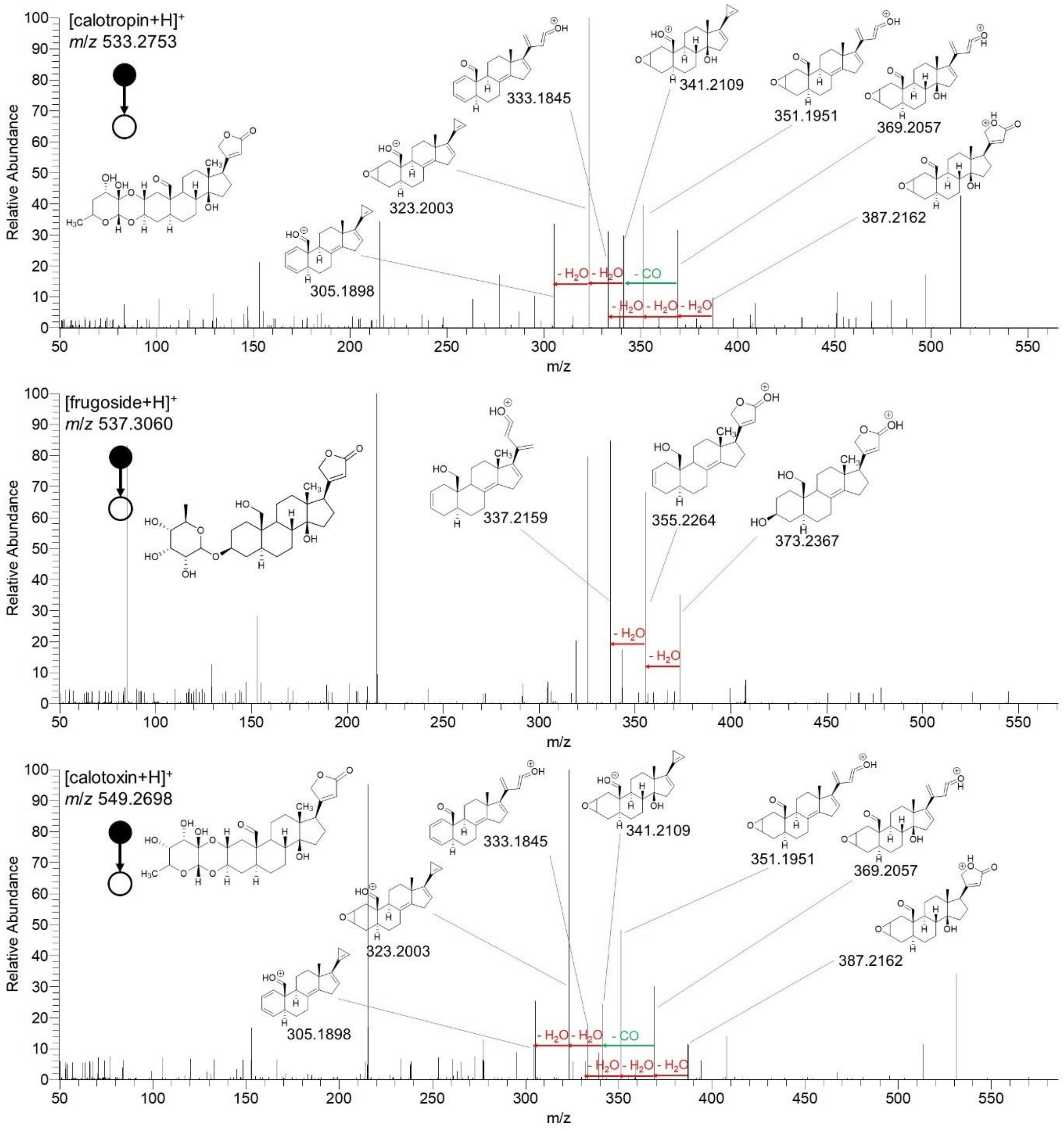


**Figure S6**. HPLC-MS/MS spectra for calotropin, frugoside and calotoxin to confirm imaging annotations. Fragmentation pathways including characteristic fragments are highlighted.

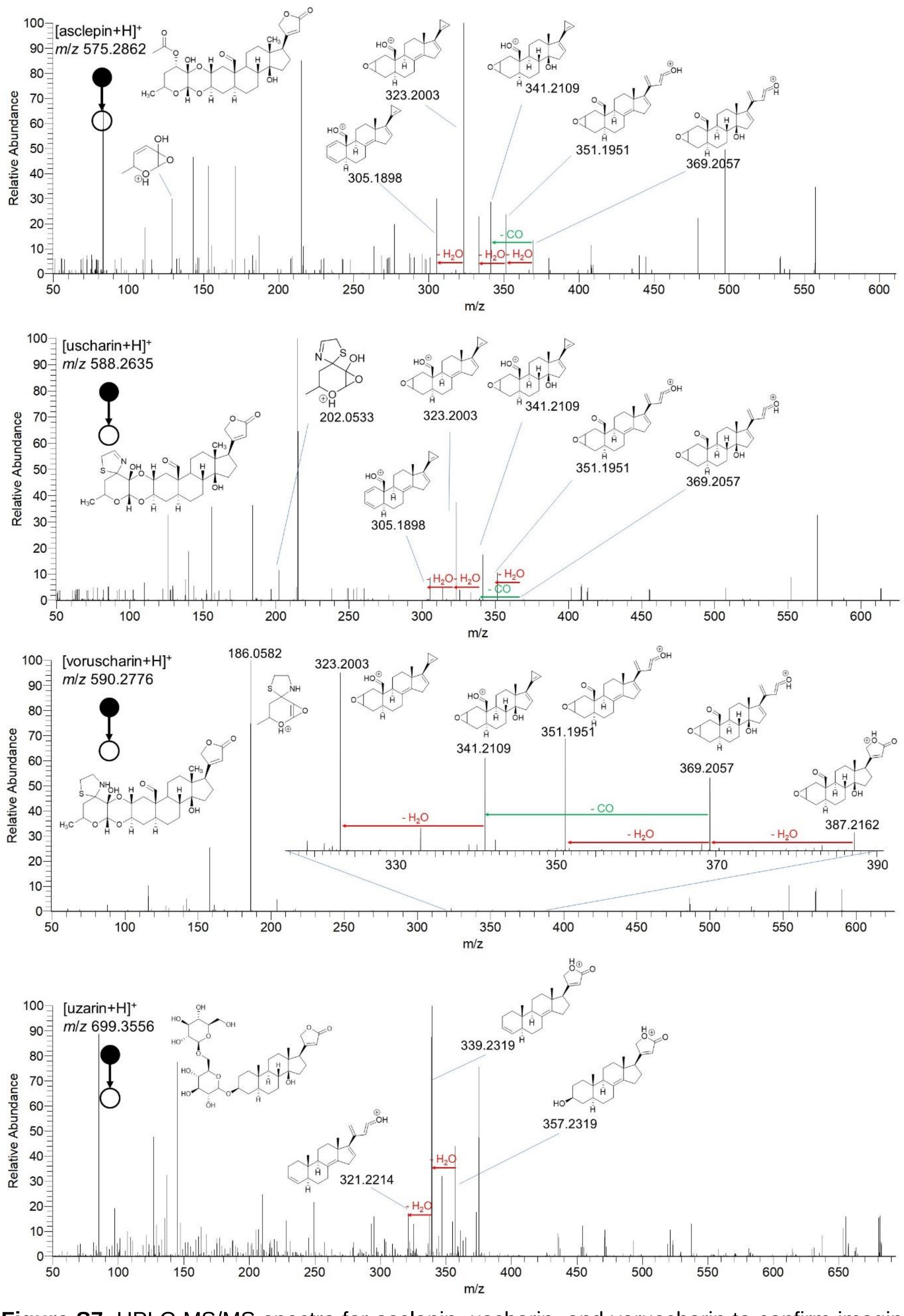


**Figure S7**. HPLC-MS/MS spectra for asclepin, uscharin, and voruscharin to confirm imaging annotations. Fragmentation pathways including characteristic fragments are highlighted.

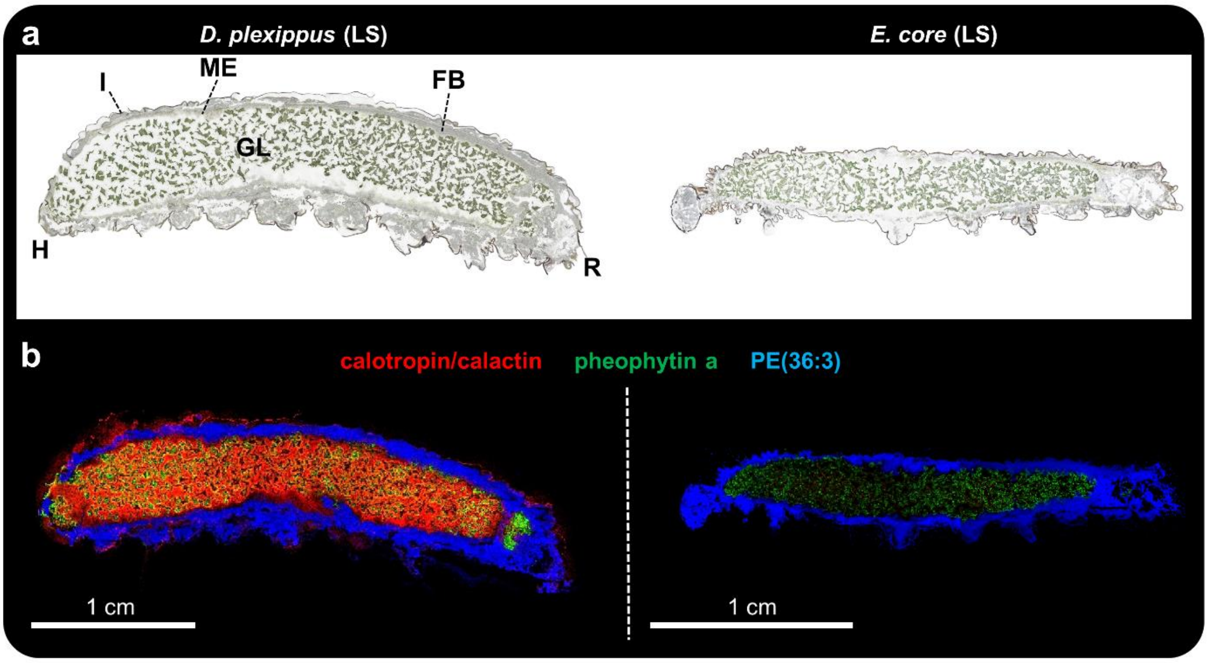


**Figure S8.** Whole-body AP-SMALDI MSI of fifth instar *D. plexippus* (biological replicate 4) and *E. core* caterpillars (biological replicate 4). (**a**) Optical image of longitudinal *D. plexippus* (left) and *E. core* (right) sections before matrix application. H: head, I: integument, R: rectum, GL: gut lumen, FB: fat body, ME: midgut epithelium. (**b**) Corresponding RGB overlay images obtained with 45 µm step size, showing the spatial distribution of calactin/calotropin ($[M+K]^+$, at *m*/*z* 571.2304 (red), pheophytin a ($[M+K]^+$, green) at *m*/*z* 909.5292 to highlight ingested *A. curassavica* plant material, and PE(36:3) ($[M+K]^+$, blue) at *m*/*z* 780.4941 serving as a marker for insect tissue, such as gut epithelium and fat body. Both RGB overlay images are normalized to the same intensity scale.

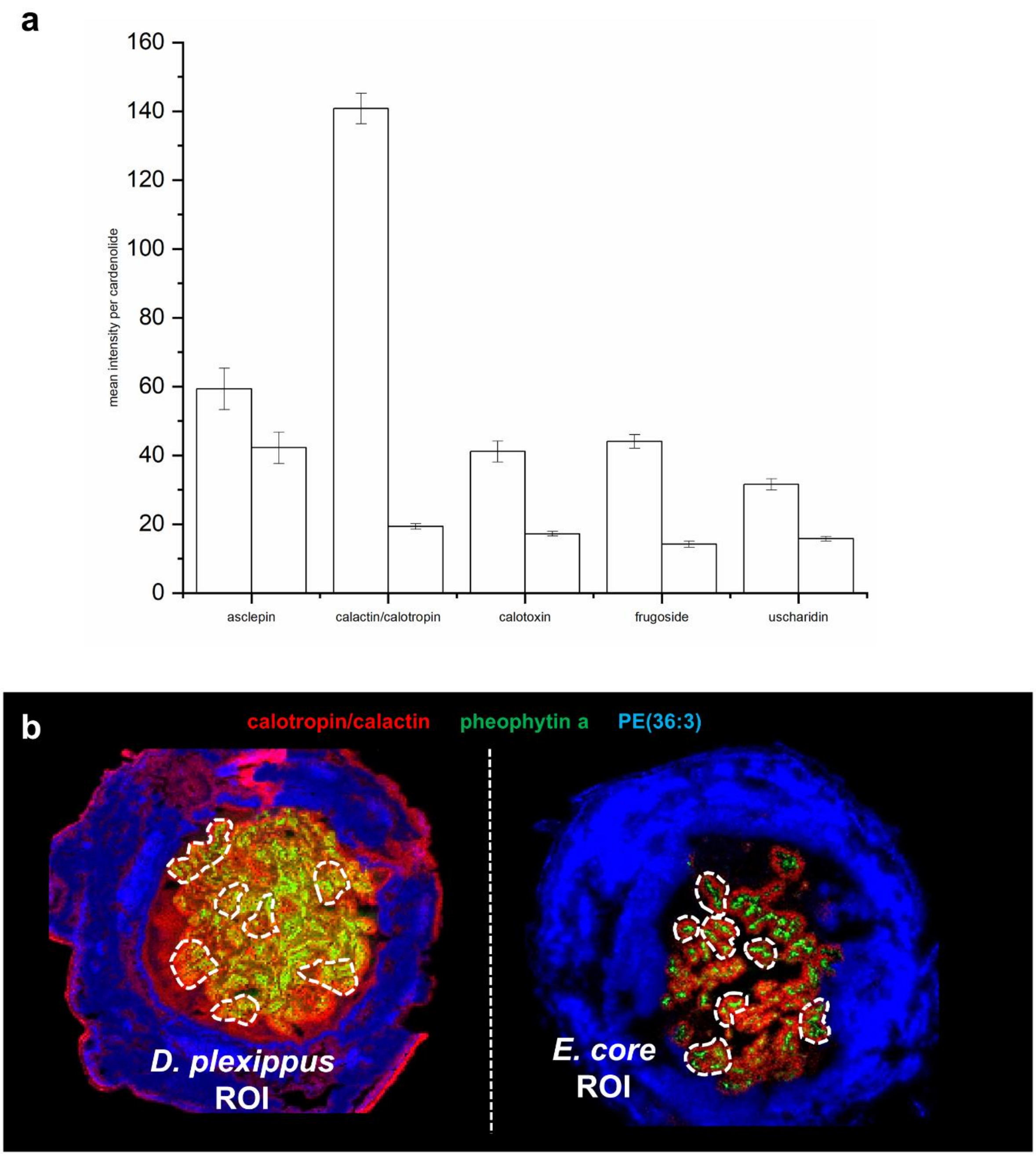


**Figure S9.** (**a**) MSI-based relative quantification for five selected cardenolide signals in the gut lumen of *D. plexippus* (left) and *E. core* (right). Bars represent means ± SE of the respective cardenolide signals based on four biological replicates (two transversal- and two longitudinal sections for each species). For each biological replicate, the mean cardenolide intensity of seven comparable regions of interest (ROI) was calculated. (**b**) Representative *in silico* ROI selection at the example of biological replicate 1 for *D. plexippus* and *E. core*. RGB overlay images showing the spatial distribution of the cardenolide calotropin and/or its isomer calactin ($[M+K]^+$, red) at *m*/*z* 571.2304, the chlorophyll derivative pheophytin a at *m*/*z* 909.5288 ($[M+K]^+$, green) as a chemical marker for plant tissue and the animal lipid PE(36:3) ($[M+K]^+$, blue) as a chemical marker for animal tissues. MSiReader was utilized to define the respective regions of interest (n = 7, total amount of pixels: 5134 pixels for *D. plexippus* and 5049 pixels for *E.*

*core*) for both species. Next, ion intensities of the respective cardenolide signal was extracted and for internal normalization and better comparison, normalized to the average pheophytin a signal abundance for the respective region of interest.

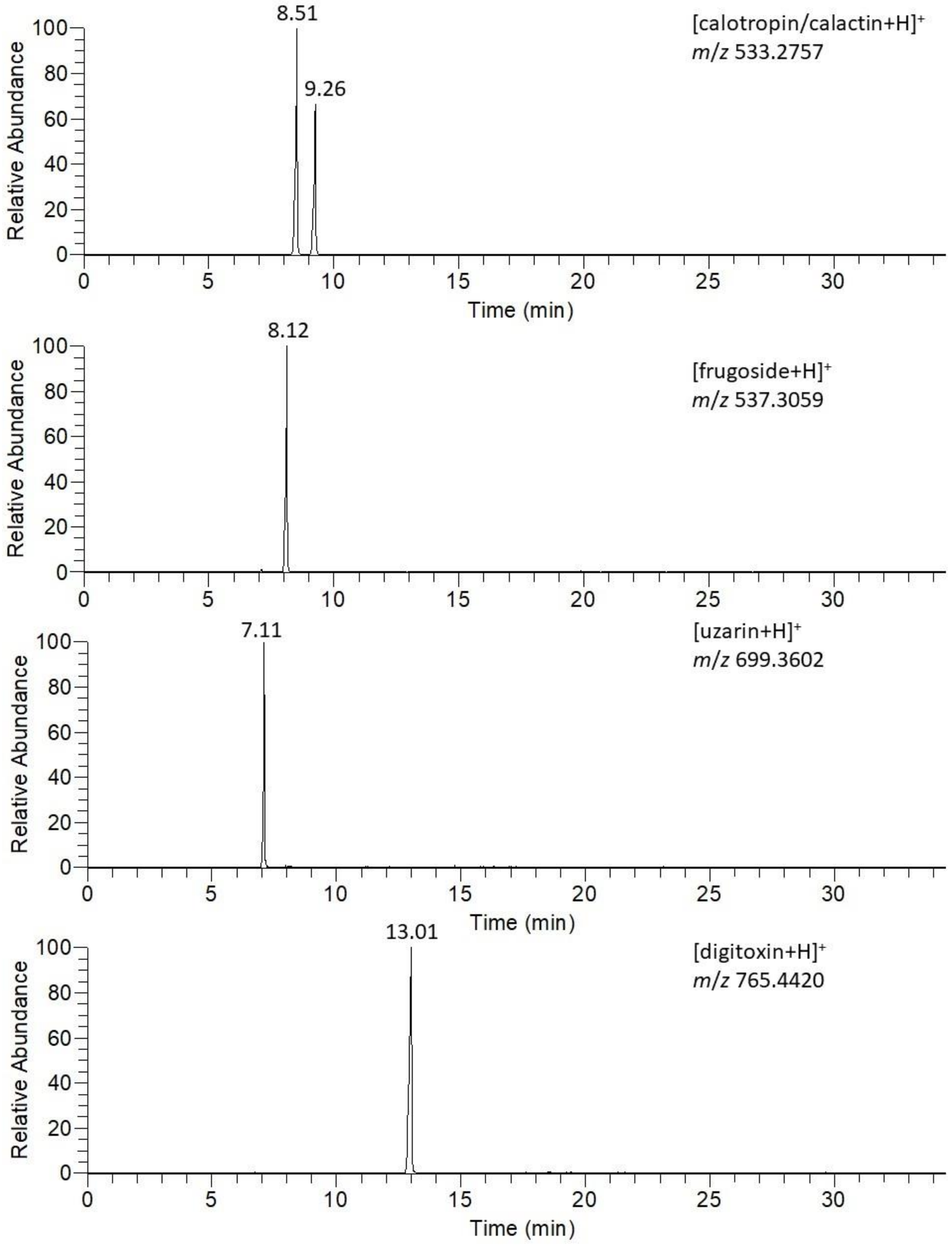


**Figure S10**. Representative HPLC-MS chromatograms of the cardenolides calotropin, calactin, frugoside, uzarin and the internal standard digitoxin, obtained from the analysis of gut material of *D. plexippus*.

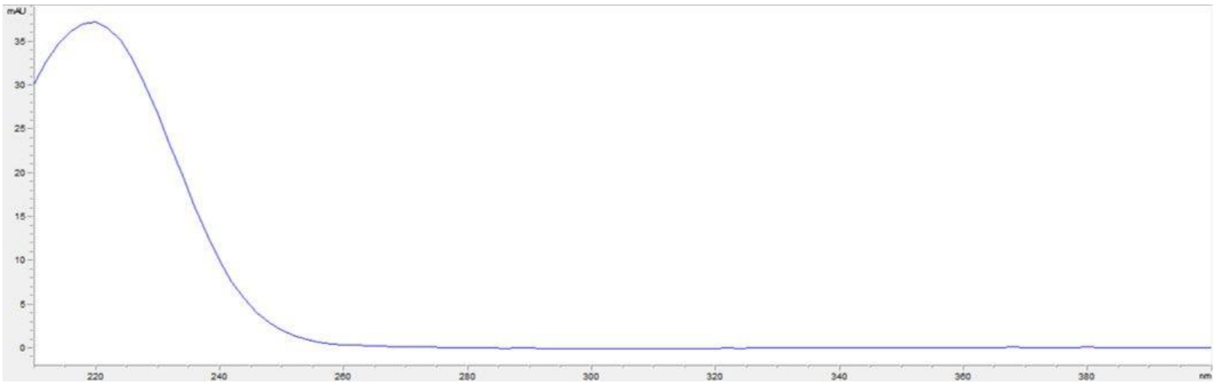

**Figure S11**. Typical DAD-spectrum of a cardenolide.

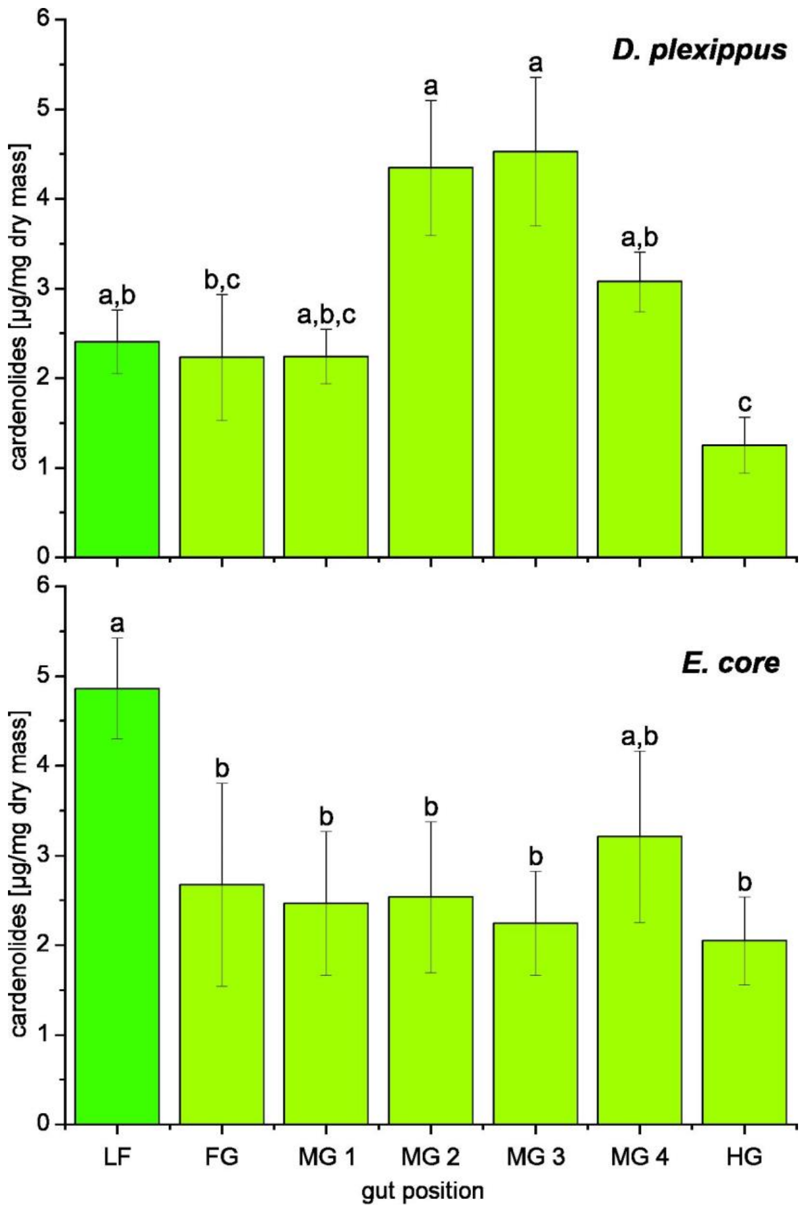


**Figure S12**. Total cardenolide concentrations across the gut passage in caterpillars of *D. plexippus* (DP) and *E. core* (EC). After dissection and freeze drying, guts were divided into foregut (FG), four portions of midgut (MG1-MG4), and hindgut (HG) and analyzed via HPLC-DAD. Samples sizes for *D. plexippus* were: n = 7 for FG, for all other gut portions n = 9; for *E. core*, sample sizes were n = 6 for all gut portions. In addition, we analyzed cardenolides in milkweed (*A. curassavica*) leaves (LF, n = 9 for DP and n = 6 for EC). Bars represent means ± SE, different letters above bars indicate statistically significant differences ($p < 0.05$). Note that cardenolide concentrations in *A. curassavica* leaves were about twice as high in the experiment with *E. core* compared to the experiment with *D. plexippus* which might be due to different environmental conditions for plant growth (greenhouse vs. climate chamber).

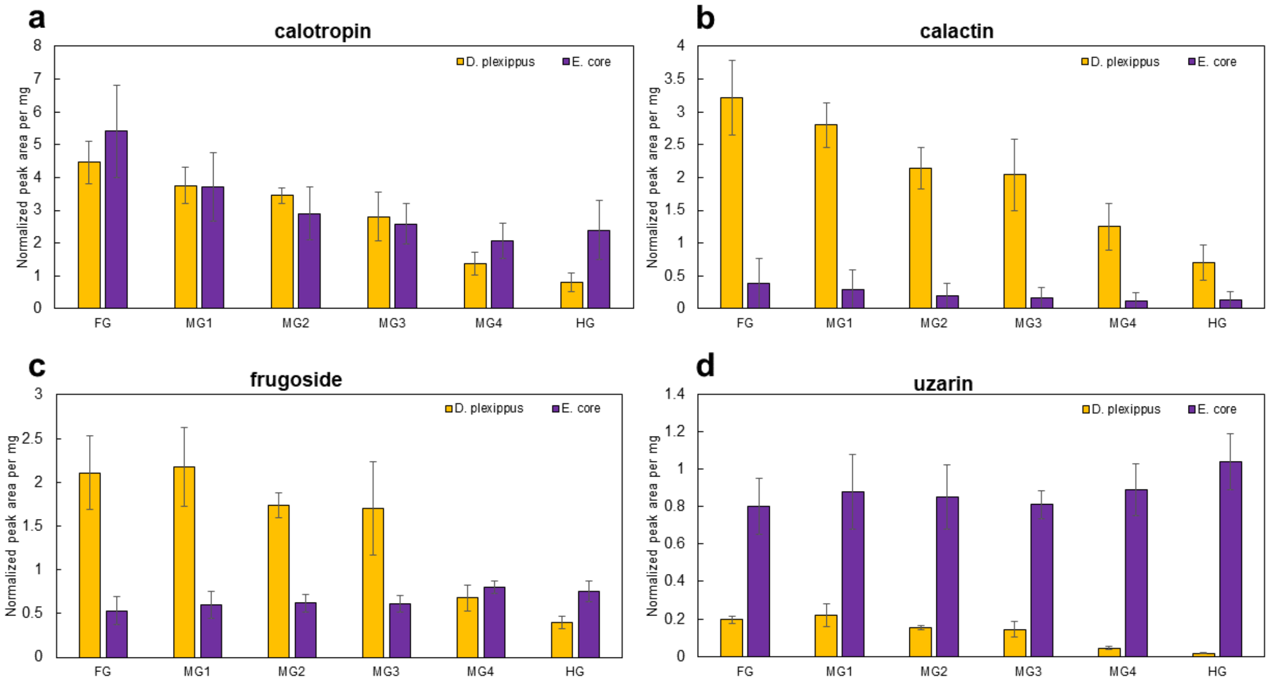


**Figure S13**. HPLC-MS-based absolute quantification of three preferentially sequestered cardenolides (calotropin, calactin, frugoside) and uzarin (not sequestered by *D. plexippus*) for gut material of *D. plexippus* (orange) and *E.core* (purple). The complete gut passage was dissected into six segments, starting with the foregut (FG), followed by 4 segments of the midgut (MG1-MG4), and the hindgut (HG). Each bar represents the mean of three biological replicates and error bars indicate standard deviation. The cardenolide peak area was normalized on the internal standard (digitoxin) and on the extracted tissue weight. (**a**) calotropin ($[M+H]^+$) at *m*/*z* 533.2755. (**b**) calactin ($[M+H]^+$) at *m*/*z* 533.2755. (**c**) frugoside ($[M+H]^+$) at *m*/*z* 537.3061. (**d**) uzarin ($[M+H]^+$) at *m*/*z* 699.3594.

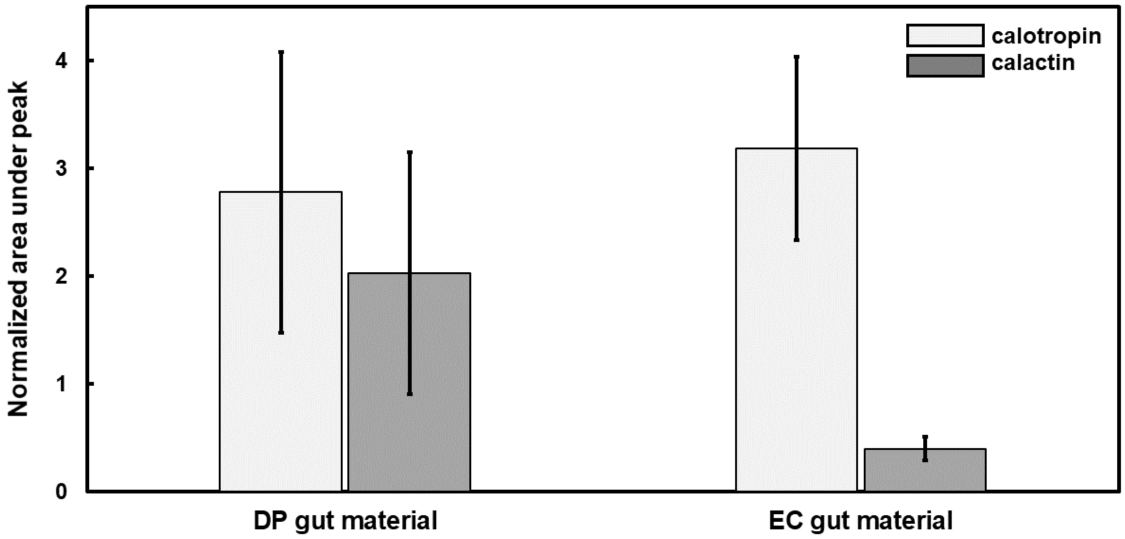


**Figure S14**. HPLC-MS-based quantification of the stereoisomers calotropin and calactin ($[M+H]^+$ at *m*/*z* 533.2751) for ingested plant material for the complete gut passage (foregut, midgut 1-4, and hindgut) of *D. plexippus* (DP) and *E. core* (EC). Each bar is the mean value of three biological replicates and error bars indicate standard deviation. The peak area was normalized on the internal standard (digitoxin) and on the extracted tissue weight of DP-/EC-gut material.

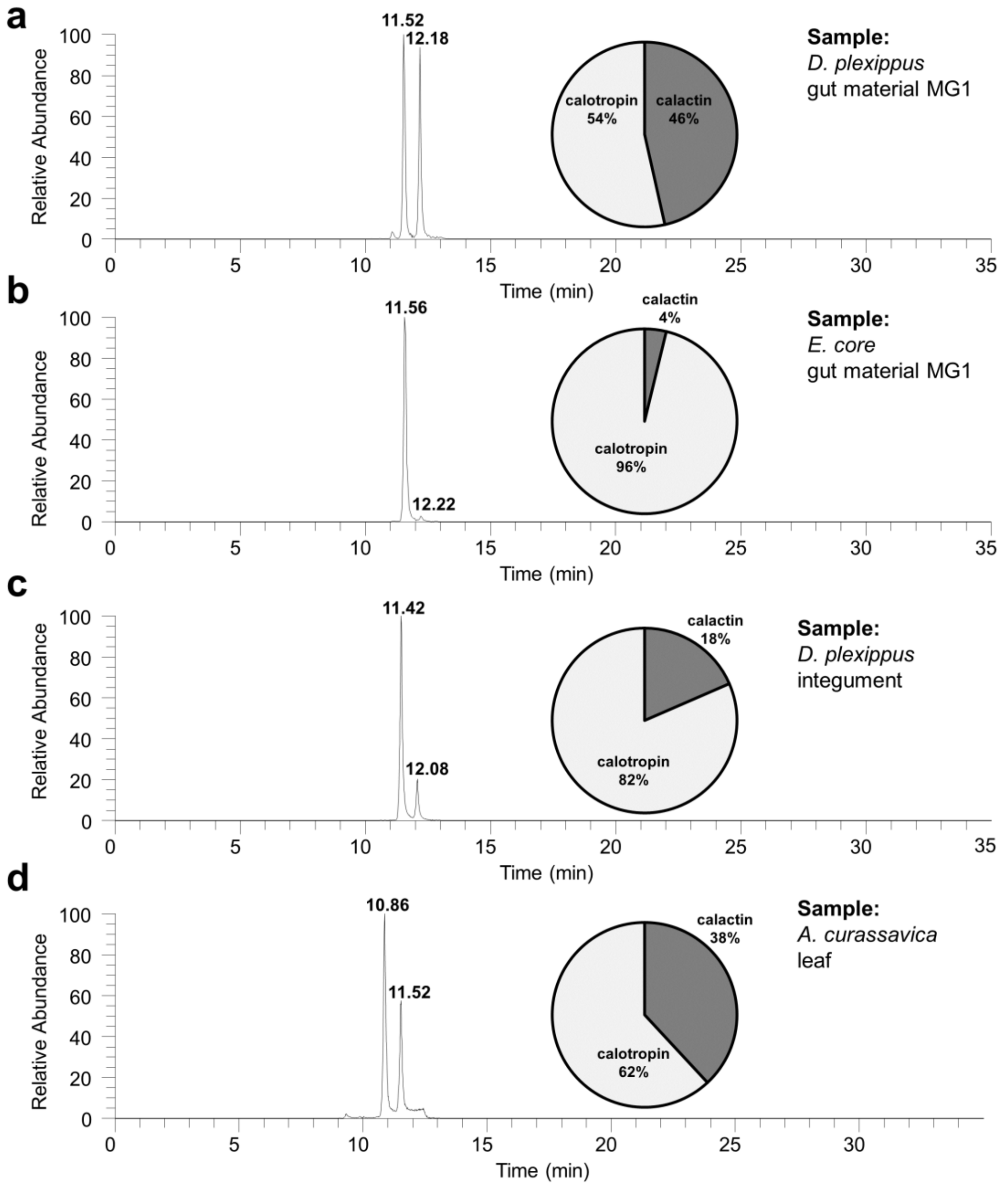


**Figure S15**. Typical HPLC-MS chromatograms for the stereoisomers calotropin and calactin ($[M+H]^+$ at *m/z* 533.2751) for (**a**) ingested plant material of *D. plexippus* midgut segment 1 (MG 1), (**b**) ingested plant material of *E. core* midgut segment 1 (MG 1), (**c**) *D. plexippus* integument tissue and (**d**) *A. curassavica* leaf tissue. The pie charts display the calotropin/calactin ratio based on the area under peak (AUP) for each sample type (n = 3).

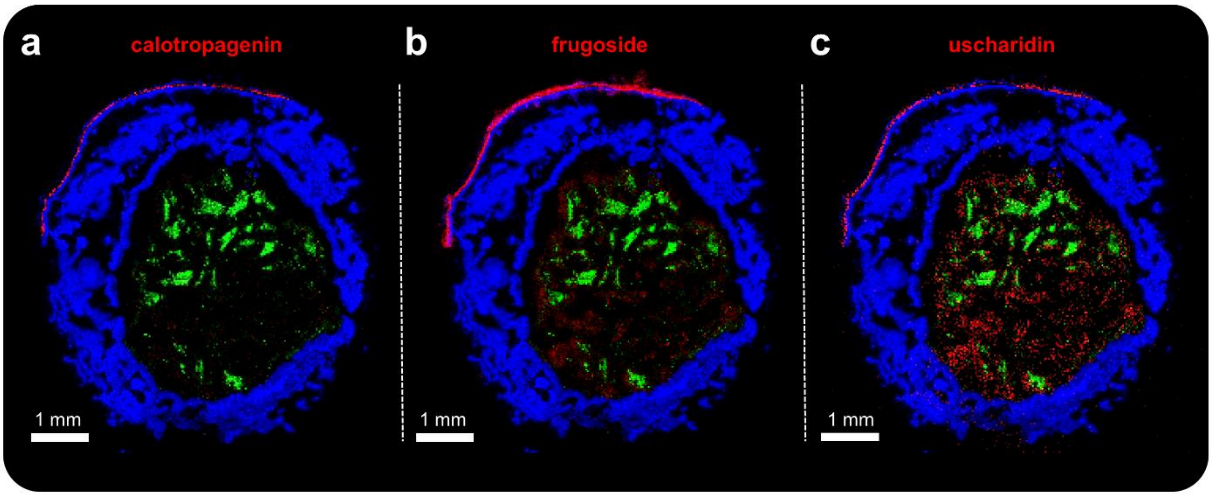


**Figure S16**. AP-SMALDI MSI (25 µm step size) of *Danaus plexippus* (biological replicate 5) fed with the non-toxic plant *Oxypetalum coeruleum* for 3 hours before sampling. (**a-c**) RGB overlay showing (**a**) calotropagenin ($[M+K]^+$, red) at *m*/*z* 443.1833, (**b**) frugoside ($[M+K]^+$, red) at *m*/*z* 575.2617, (**c**) uscharidin ($[M+K]^+$, red) at *m*/*z* 569.2146, and (**a-c**) pheophytin a ($[M+K]^+$, green) at *m*/*z* 909.5292 and PE(36:3) ($[M+K]^+$, blue) at *m*/*z* 780.4942.

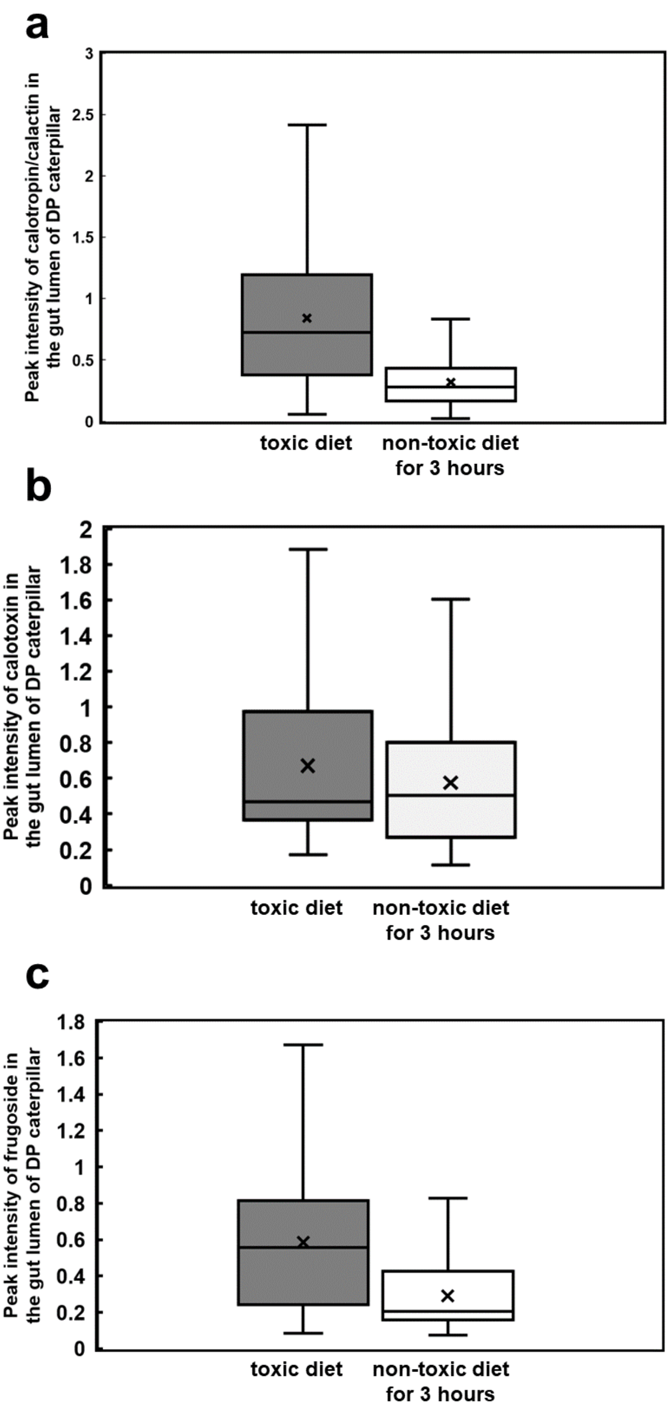


**Figure S17**. MSI-based relative quantification for the preferentially-sequestered cardenolides in the gut lumen of transversally sectioned caterpillars of *Danaus plexippus*. The respective cardenolide abundance was compared between caterpillars fed with *Asclepias curassavica* (deep grey, MSI data from Fig. 1, biological replicate 1) diet and caterpillars raised on *A. curassavica* but purged with a cardenolide-free diet of *Oxypetalum coeruleum* (light grey, MSI data from Fig. 3, biological replicate 5) for 3 hours before sampling. MSiReader was utilized to *in silico* segment the gut lumen and integument and subsequently, cardenolide signal intensities were extracted. Previous LC-MS-based cardenolide quantification for *D. plexippus* integument tissue (Fig. S19) showed no significant variations across biological replicates. For better comparison, cardenolide signal intensities from the gut lumen were normalized to the average signal abundance of the respective cardenolide at the integument for the same MSI experiment. Box plots for (**a**) calotropin/calactin ($[M+K]^+$ at *m*/*z* 571.2304) showing 2.7 x less abundance after purging the gut passage with cardenolide-free plant material, (**b**) calotoxin ($[M+K]^+$ at *m*/*z* 587.2251) showing 1.1 x less abundance after purging the gut passage with cardenolide-free plant material, (**c**) frugoside ($[M+K]^+$ at *m*/*z* 575.2622) showing 2.1 x less abundance after purging the gut passage with cardenolide-free plant material.

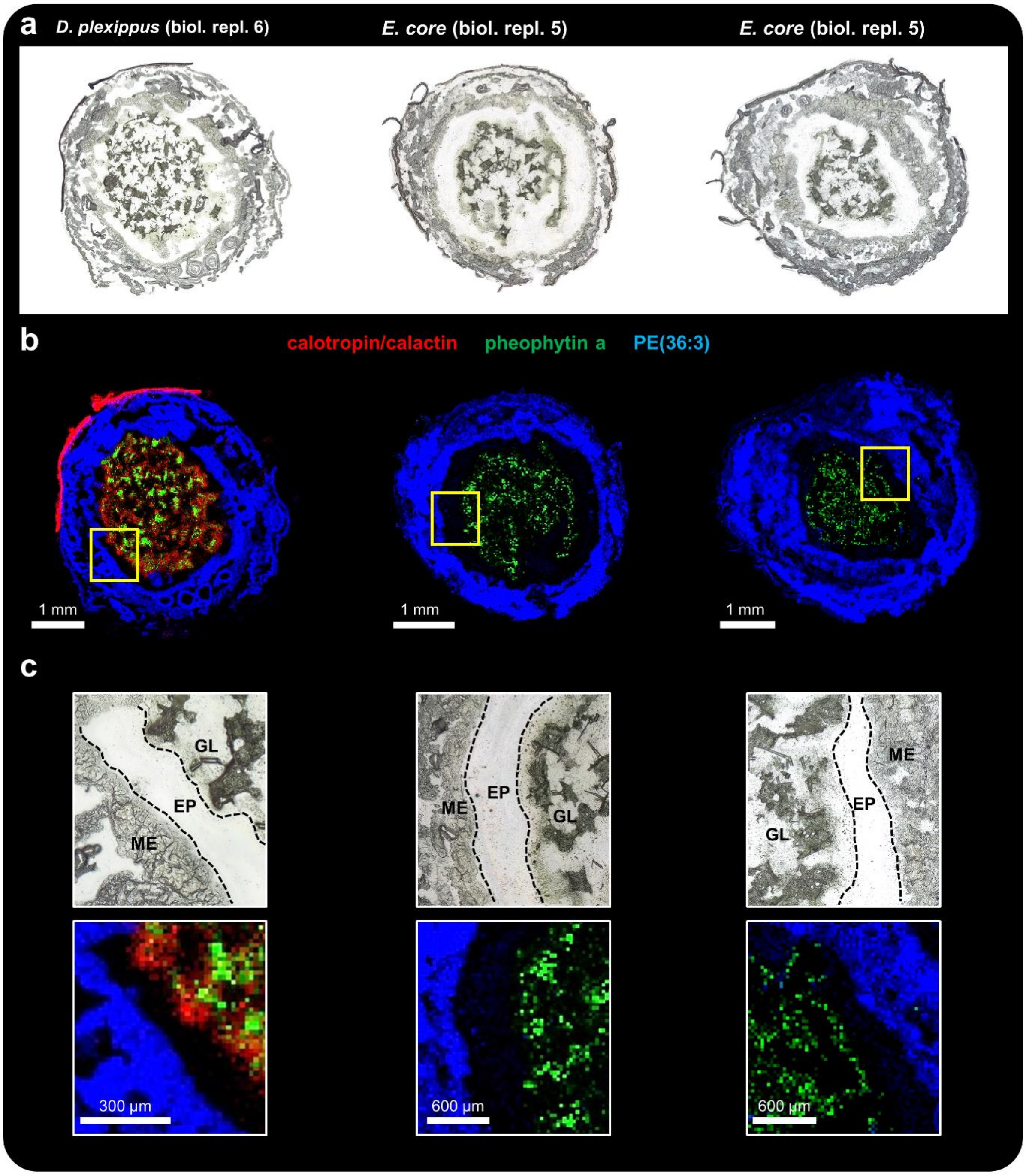


**Figure S18.** AP-SMALDI MSI (25 µm step size) of *Danaus plexippus* (biological replicate 6) and *Euploea core* caterpillars (biological replicate 5) purged with the non-toxic plant *Oxypetalum coeruleum* before sampling. (**a**) Optical images of transversally sectioned last instar *D. plexippus* and *E. core* caterpillars. (**b**) Corresponding RGB overlay images showing calactin/calotropin ($[M+K]^+$, red) at *m*/*z* 571.2304, pheophytin a ($[M+K]^+$, green) at *m*/*z* 909.5292 and PE(36:3) ($[M+K]^+$, blue) at *m*/*z* 780.4942. (**c**) Magnified views of the outlined regions in **b** highlighting midgut epithelium tissue (ME), ectoperitrophic space (EP) and gut lumen (GL).

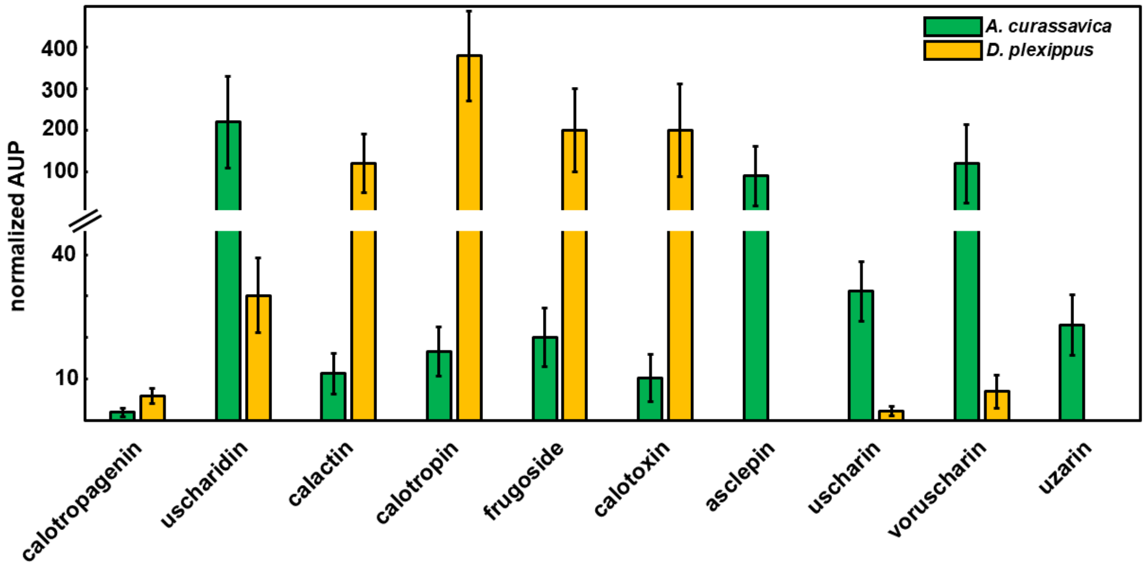


**Figure S19.** HPLC-MS-based quantification for 10 detected cardenolides in *Asclepias curassavica* leaf material and *D. plexippus* integument. Each bar represents the mean of three biological replicates and error bars indicate standard deviation. The area under peak (AUP) was normalized to the internal standard (digitoxin). Please note that quantities between leaf material and caterpillar tissue may not be compared in absolute terms since sample dry masses were different.

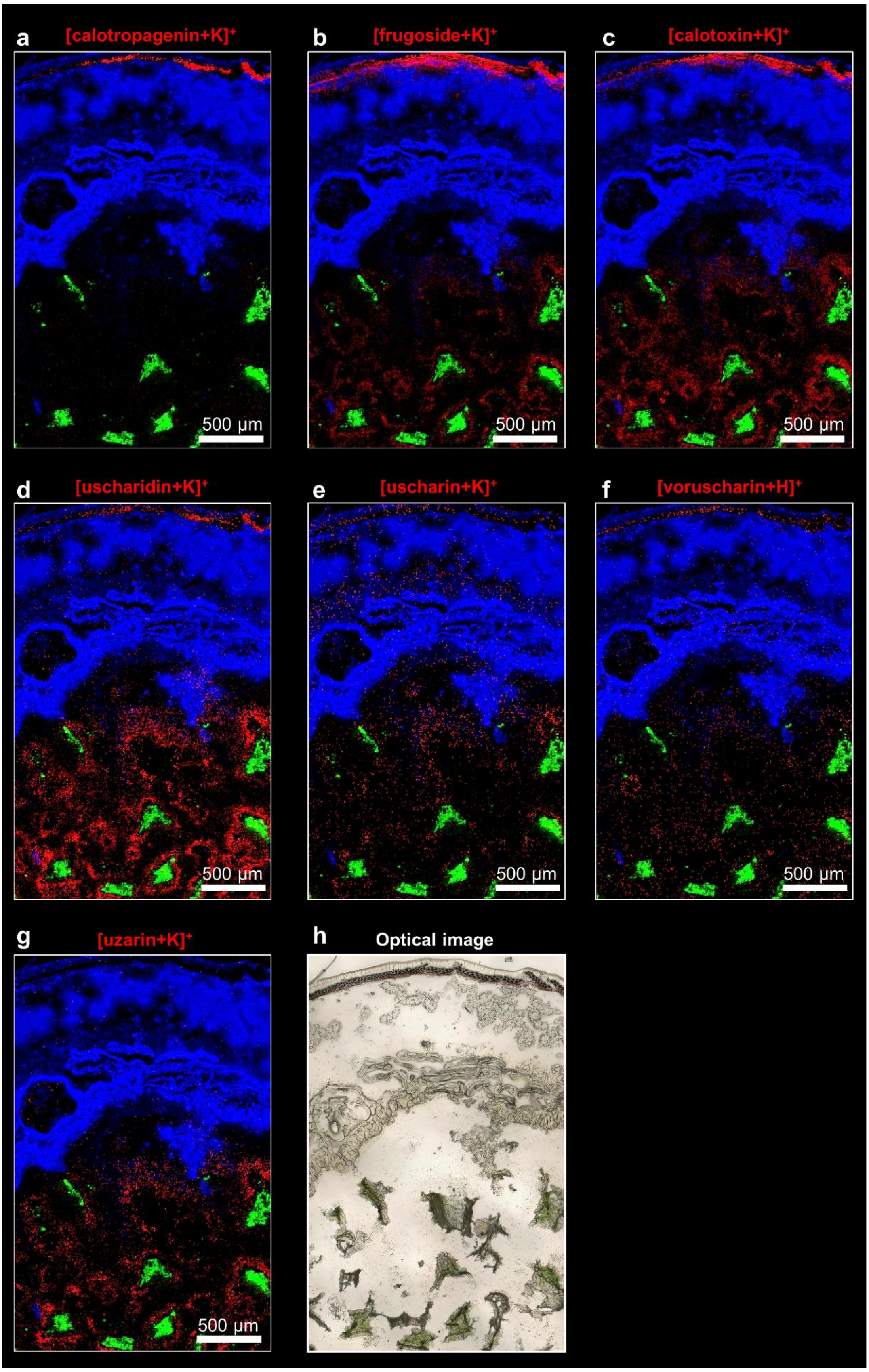


**Figure S20.** High-resolution AP-SMALDI MSI (5 µm step size) of transversal *D. plexippus* (biological replicate 7) section. RGB images showing the spatial distribution of (**a**) calotropagenin ($[M+K]^+$, red) at *m*/*z* 443.1829, (**b**) frugoside ($[M+K]^+$, red) at *m*/*z* 575.2616, (**c**)

calotoxin ([M+K]$^+$, red) at *m*/*z* 587.2253, (**d**) uscharidin ([M+K]$^+$, red) at *m*/*z* 569.2146, (**e**) uscharin ([M+K]$^+$, red) at *m*/*z* 626.2159, (**f**) voruscharin ([M+H]$^+$, red) at *m*/*z* 590.2770, (**g**) uzarin ([M+K]$^+$, red) at *m*/*z* 737.3149. Additionally, (**a-g**) pheophytin a ([M+K]$^+$, green) at *m*/*z* 909.5290 and PE(36:3) ([M+K]$^+$, blue) at *m*/*z* 780.4940 were used to highlight ingested plant tissue and insect morphology. (**h**) Optical image of the analyzed region of interest.

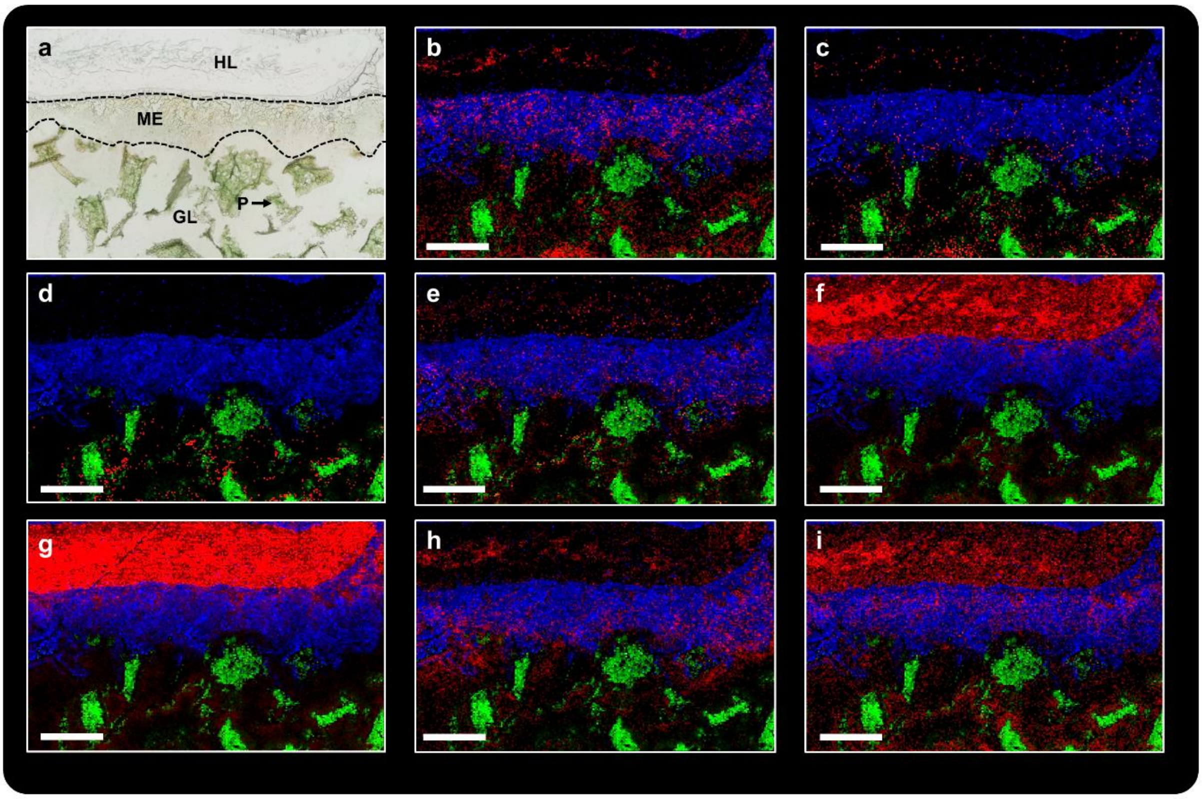


**Figure S21**. High-resolution AP-SMALDI MSI (10 µm step size) of midgut epithelium tissue in a longitudinal *Danaus plexippus* (biological replicate 3) section to investigate transport of selected cardenolide and other primary- and secondary metabolites. (**a**) Optical image of the analyzed region of interest. P: *Asclepias curassavica* plant material, GL: gut lumen, ME: midgut epithelium tissue, HL: hemolymph. (**b-i**) RGB overlay images showing the spatial distribution for (**b**) frugoside ($[M+K]^+$, red) at *m*/*z* 575.2616, (**c**) calotoxin ($[M+K]^+$, red) at *m*/*z* 587.2253, (**d**) asclepin ($[M+K]^+$, red) at *m*/*z* 613.2415, (**e**) kaempferol-glucopyranoside ($[M+Na]^+$, red) at *m*/*z* 471.0898, (**f**) malvidin-glucoside ($[M+Na]^+$, red) at *m*/*z* 515.1160, (**g**) N-(1-deoxy-1-fructosyl)tyrosine ($[M+K]^+$, red) at *m*/*z* 382.0899, (**h**) guanosine ($[M+K]^+$, red) at *m*/*z* 322.0548, (**i**) disaccharide ($[M+K]^+$, red) at *m*/*z* 381.0793. Additionally, (**b-i**) pheophytin a ($[M+K]^+$, green) at *m*/*z* 909.5290 and PE(36:3) ($[M+K]^+$, blue) at *m*/*z* 780.4940 were used to highlight ingested plant tissue and midgut epithelium tissue. Scale bars, 500 µm.

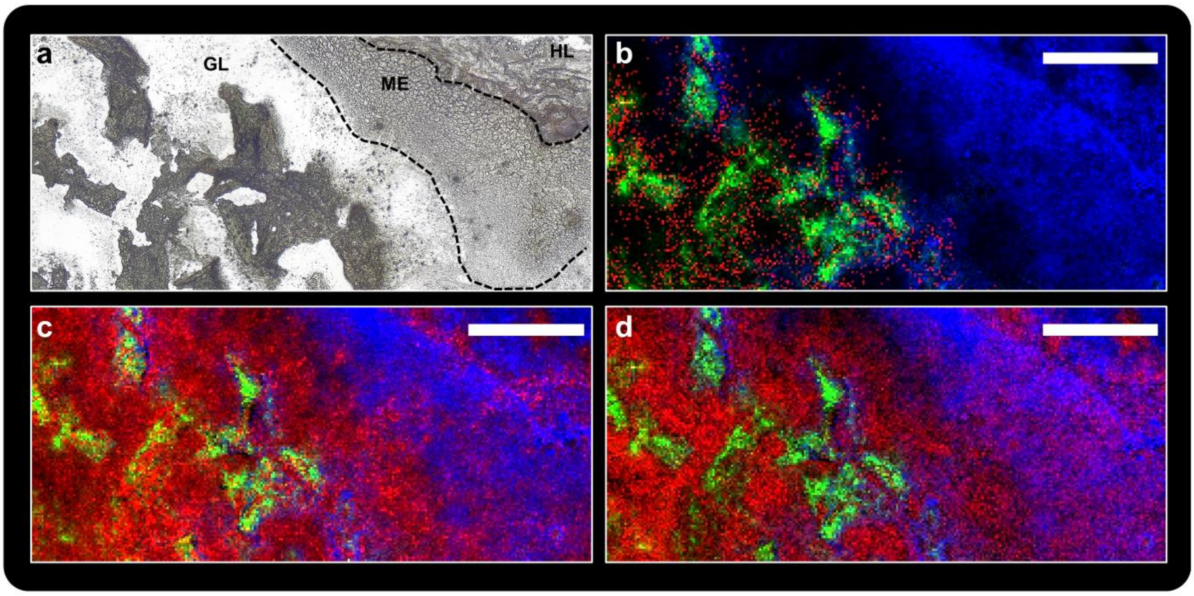


**Figure S22**. High-resolution AP-SMALDI MSI (10 µm step size) of midgut epithelium tissue of a longitudinal *Euploea core* (biological replicate 3) section to investigate various primary- and secondary metabolites. (**a**) Optical image of the analyzed region of interest. GL: gut lumen, ME: midgut epithelium tissue, HL: hemolymph. (**b**-**d**) RGB overlay images showing the spatial distribution for (**b**) calotropin/calactin ($[M+K]^+$, red) at *m*/*z* 571.2305, (**c**) guanosine ($[M+K]^+$, red) at *m*/*z* 322.0547, (**d**) disaccharide ($[M+K]^+$, red) at *m*/*z* 381.0793 and (**b-d**) pheophytin a ($[M+K]^+$, green) at *m*/*z* 909.5290 and PE(36:3) ($[M+K]^+$, blue) at *m*/*z* 780.4940. Scale bars, 500 µm.

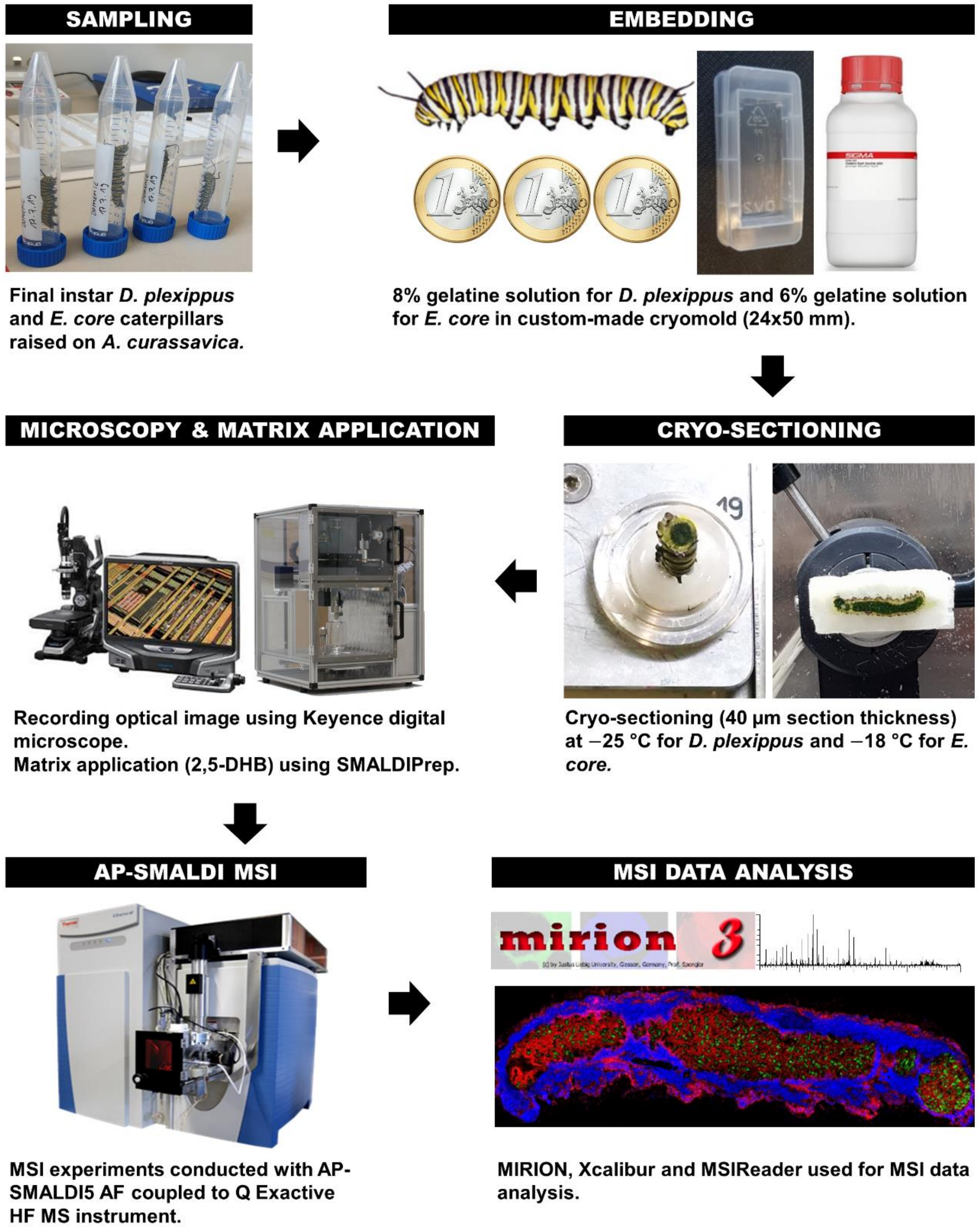


**Figure S23**. MSI-based spatial metabolomics workflow to study the distribution of sequestered cardenolides in milkweed butterflies.

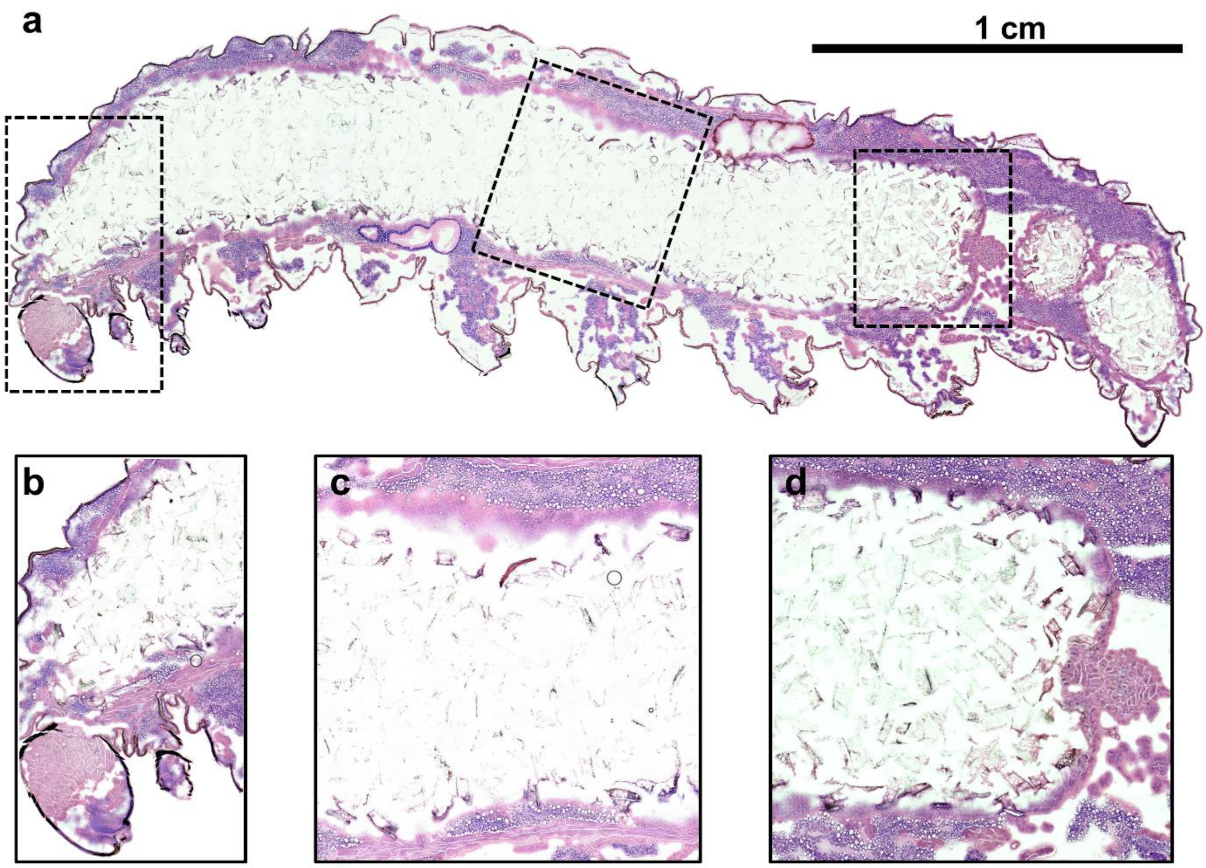


**Figure S24**. Optical image of a longitudinal monarch section (biological replicate 3), which was analysed via AP-SMALDI MSI (see Figure 2). The morphology and different tissue types remain unperturbed during AP-SMALDI MSI analysis, thus allowing to perform classical histology on the same section. The matrix layer was removed with 70% ethanol and H&E-staining was carried out. (**a**) Whole-body of the final instar caterpillar, (**b**) anterior region showing mid gut epithelium tissue, (**c**) central region of the mid gut, (**d**) end of the mid gut.

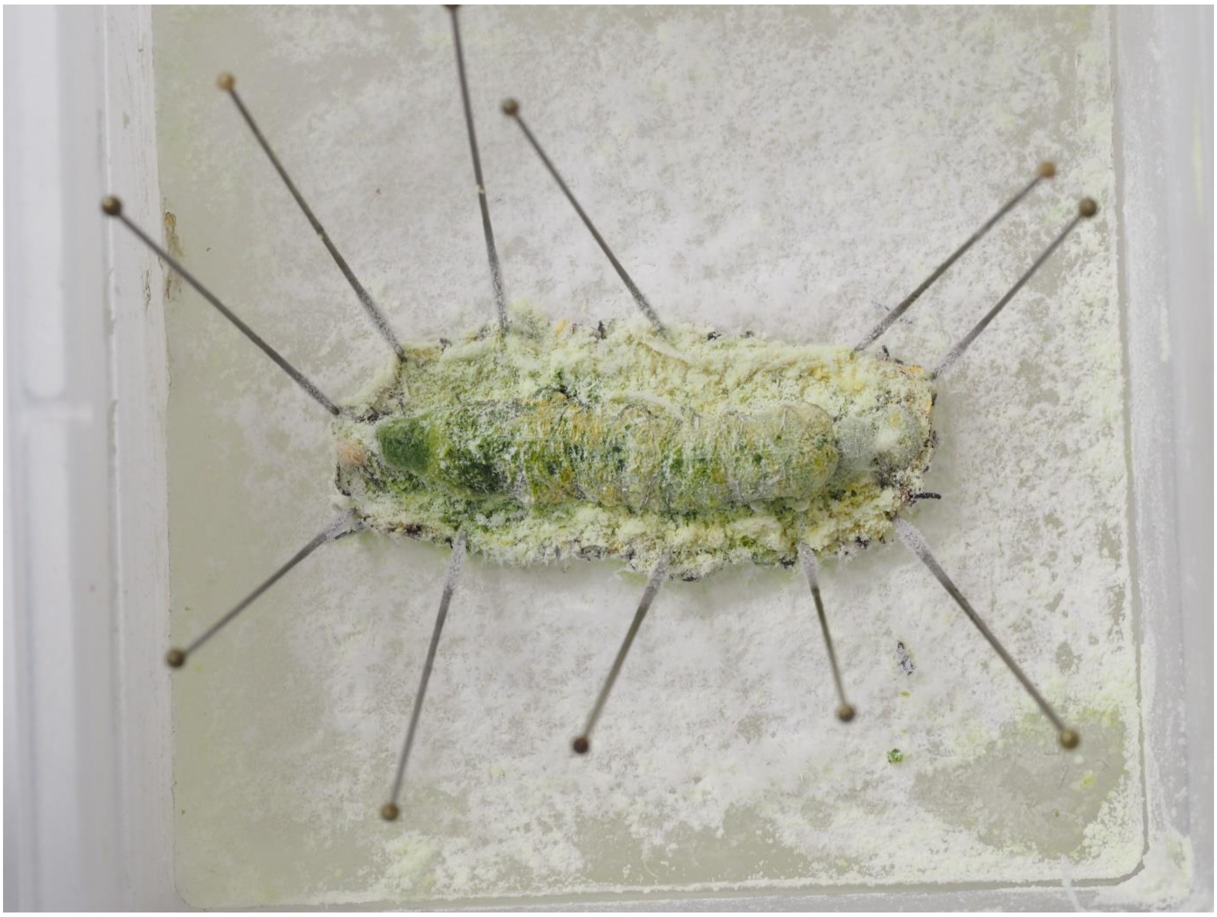

**Figure S25**. *Danaus plexippus* caterpillar freeze dried after dissection. Precipitated salts were removed using a soft brush before the gut was divided into foregut (left), four portions of midgut, and hindgut (right). Please see the methods section of the manuscript for further details.

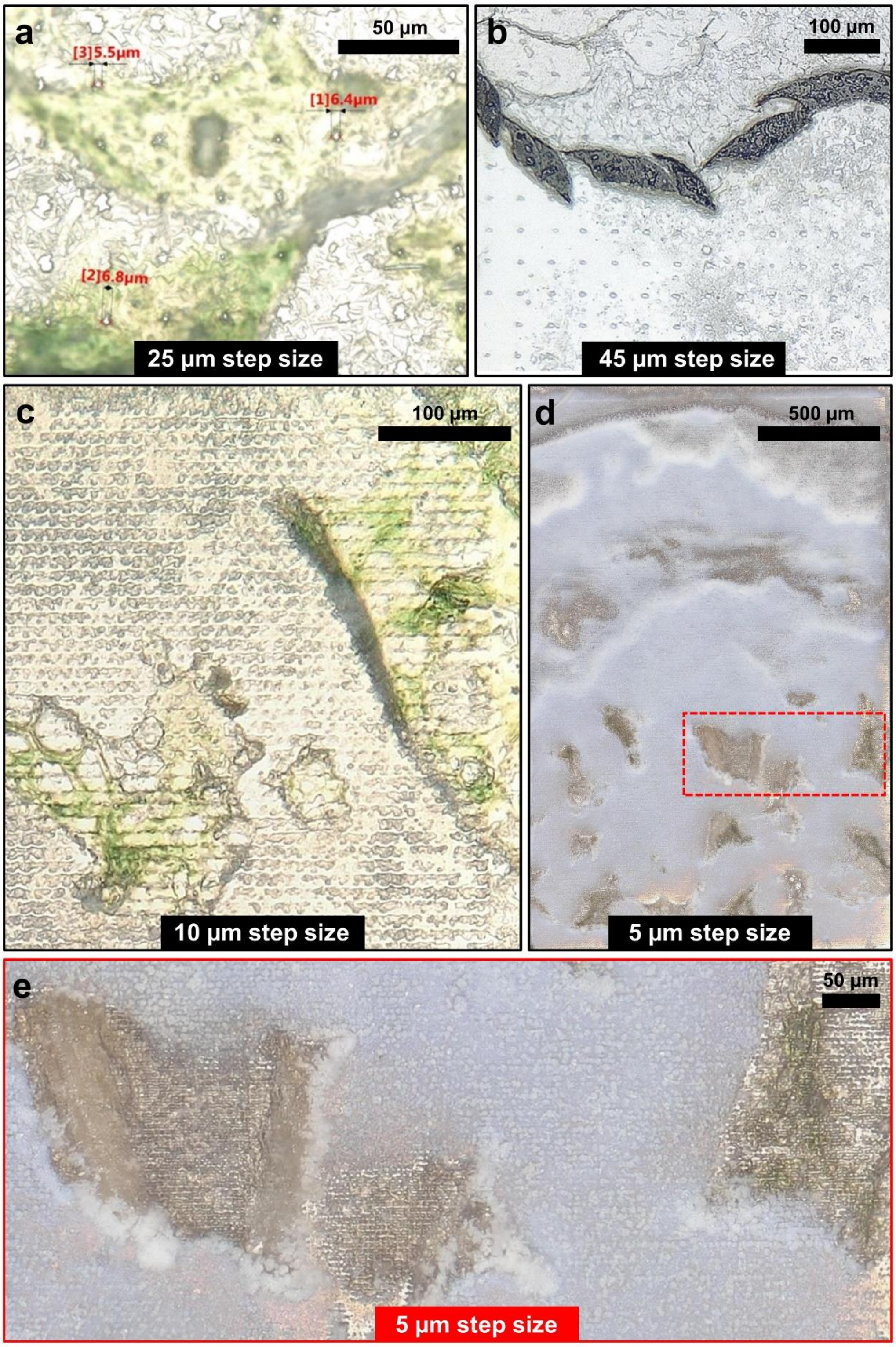


**Figure S26**. Laser ablation spots in the crystalline matrix layer (2,5-DHB) of *D. plexippus* tissue sections for AP-SMALDI MSI experiments conducted with (**a**) 25 µm, (**b**) 45 µm, (**c**) 10 µm and (**d,e**) 5 µm step-size. The laser settings were carefully adjusted in respect to the applied step-size to prevent oversampling (i.e. shooting the same sampling area twice). Of especial importance for low-micrometer-resolution MSI (step-sizes below 10 µm) is that the employed AP-SMALDI5 AF ion source provides a laser focus of 5 µm, thus, allowing us to perform the respective MSI experiments without oversampling.

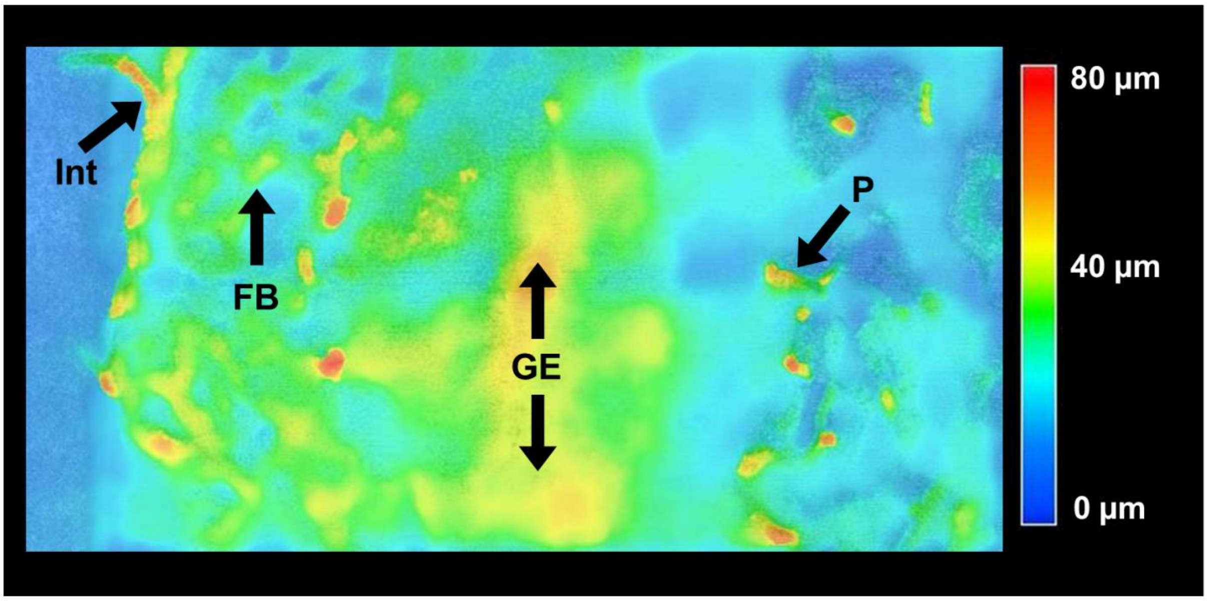


**Figure S27**. Topography image of a transversal *E. core* section showing height variations of up to 80 µm at ingested *A. curassavica* plant material (P) and integument (Int). Across midgut epithelium (GE) and fat body (FB), we found height variations of up to 40 µm. Therefore, we conducted high-resolution MSI experiments (5 µm to 10 µm step size) using the 3D-surface mode (pixelwise autofocusing) of the AP-SMALDI5 AF ion source to keep the MALDI laser focus, fluence and ablation spot size constant for the whole MSI experiment.

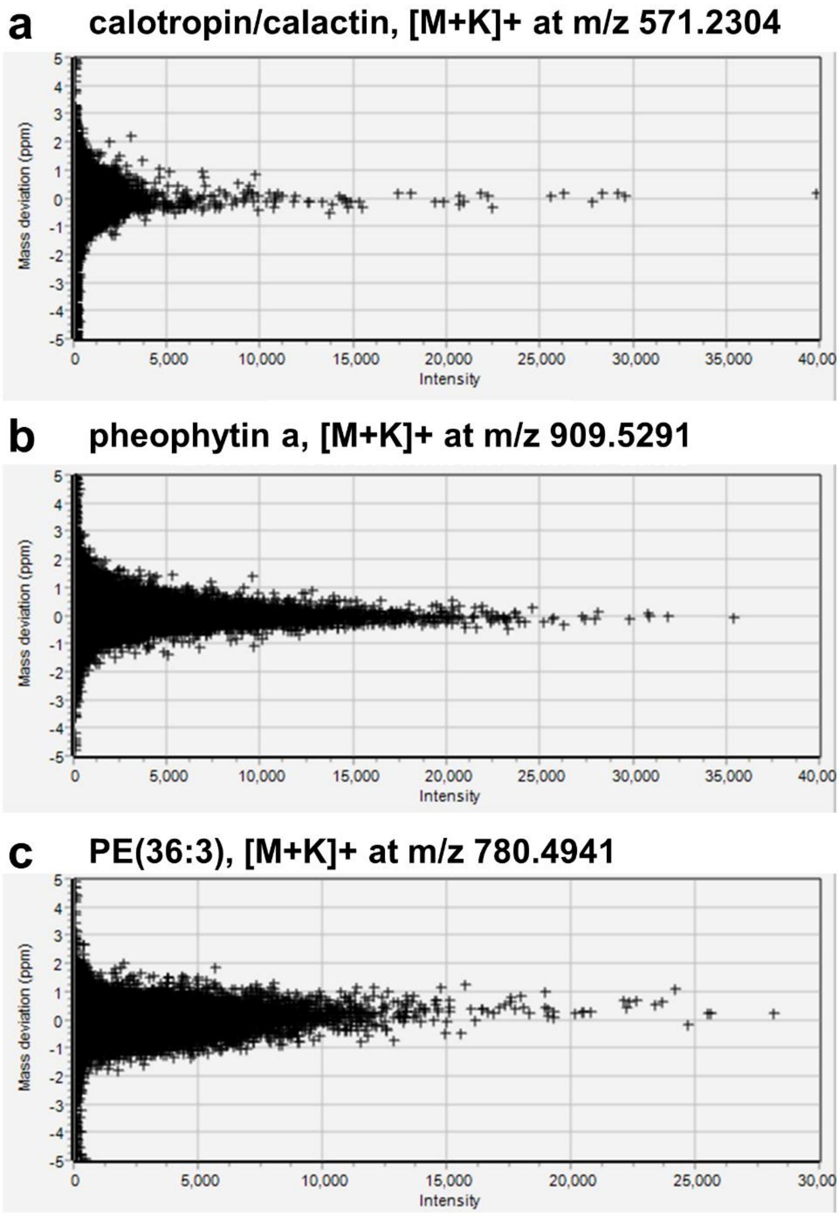


**Figure S28**. Root-mean-square error plots for (**a**) calotropin/calactin ([M+K]+ at *m*/*z* 571.2304), (**b**) pheophytin a ([M+K]+ at *m*/*z* 909.5291) and (**c**) PE(36:3) ([M+K]+ at *m*/*z* 780.4941).

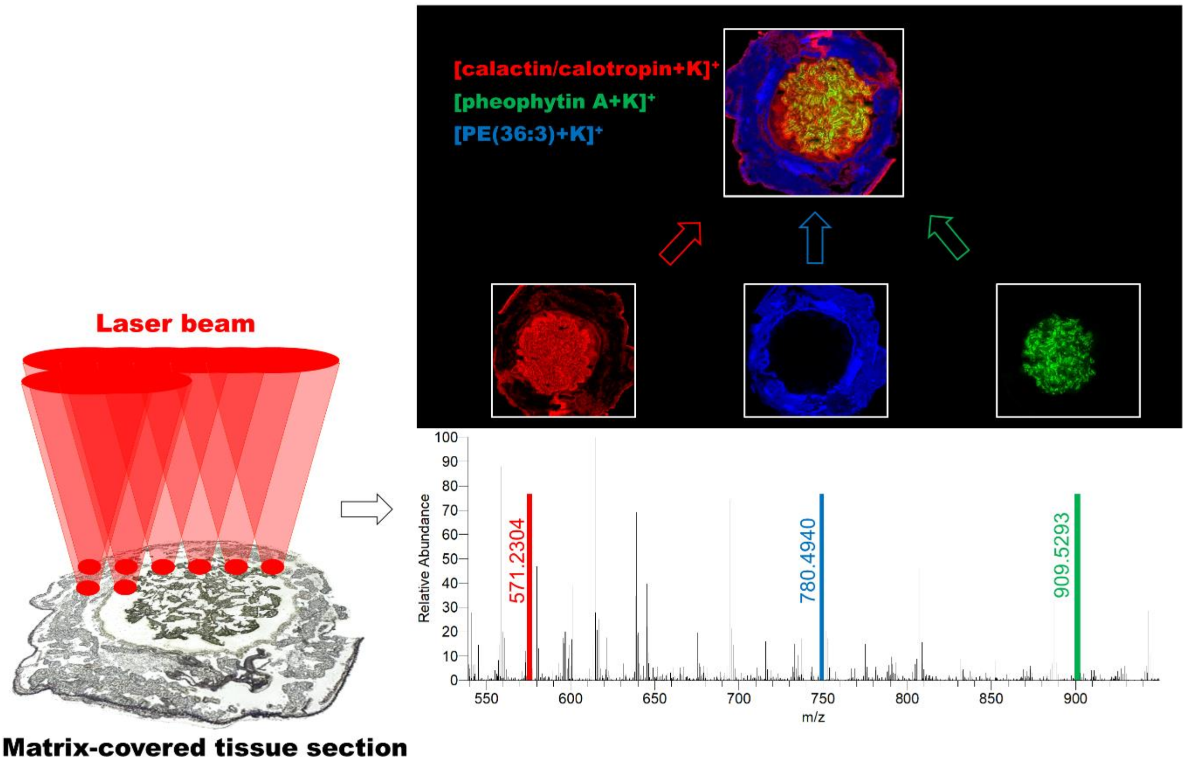


**Figure S29**. General principle to obtain red-green-blue (RGB) overlay ion images using AP-SMALDI MSI. The matrix-covered sample tissue section is scanned by a laser beam for desorption/ionization of analytes with a defined step size between each measurement spot. For image generation, the intensity values of the selected mass-to-charge-number ratio (*m*/*z*) are combined with the spatial information into an image which displays the spatial distribution of the selected signal on the sample surface. For the RGB overlay image, three different signals are assigned to one colour each and superimposed.
